# Metastatic site influences driver gene function in pancreatic cancer

**DOI:** 10.1101/2024.03.17.585402

**Authors:** Kaloyan M. Tsanov, Francisco M. Barriga, Yu-Jui Ho, Direna Alonso-Curbelo, Geulah Livshits, Richard P. Koche, Timour Baslan, Janelle Simon, Sha Tian, Alexandra N. Wuest, Wei Luan, John E. Wilkinson, Ignas Masilionis, Nevenka Dimitrova, Christine A. Iacobuzio-Donahue, Ronan Chaligné, Dana Pe’er, Joan Massagué, Scott W. Lowe

## Abstract

Driver gene mutations can increase the metastatic potential of the primary tumor^1–3^, but their role in sustaining tumor growth at metastatic sites is poorly understood. A paradigm of such mutations is inactivation of *SMAD4* – a transcriptional effector of TGFβ signaling – which is a hallmark of multiple gastrointestinal malignancies^4,5^. *SMAD4* inactivation mediates TGFβ’s remarkable anti-to pro-tumorigenic switch during cancer progression and can thus influence both tumor initiation and metastasis^6–14^. To determine whether metastatic tumors remain dependent on *SMAD4* inactivation, we developed a mouse model of pancreatic ductal adenocarcinoma (PDAC) that enables *Smad4* depletion in the pre-malignant pancreas and subsequent *Smad4* reactivation in established metastases. As expected, *Smad4* inactivation facilitated the formation of primary tumors that eventually colonized the liver and lungs. By contrast, *Smad4* reactivation in metastatic disease had strikingly opposite effects depending on the tumor’s organ of residence: suppression of liver metastases and promotion of lung metastases. Integrative multiomic analysis revealed organ-specific differences in the tumor cells’ epigenomic state, whereby the liver and lungs harbored chromatin programs respectively dominated by the KLF and RUNX developmental transcription factors, with *Klf4* depletion being sufficient to reverse *Smad4*’s tumor-suppressive activity in liver metastases. Our results show how epigenetic states favored by the organ of residence can influence the function of driver genes in metastatic tumors. This organ-specific gene–chromatin interplay invites consideration of anatomical site in the interpretation of tumor genetics, with implications for the therapeutic targeting of metastatic disease.

## MAIN

Metastatic disease – the growth of cancers beyond the primary tumor – accounts for 90% of cancer-related deaths^15,16^. Metastasis involves the acquisition of multiple traits that enable cells to leave the primary tumor, survive in the circulation, and ultimately reach and colonize other organs^16,17^. Despite the distinct capabilities that must be acquired for a tumor cell to successfully metastasize, genome-sequencing efforts have identified few driver gene mutations that are specific to metastatic tumors^5,18,19^. This has suggested that pro-metastatic traits arise from epigenetic programs that facilitate cell state changes such as the epithelial-to-mesenchymal transitions (EMTs)^16,17,20–22^. Although driver gene mutations can endow primary tumors with increased metastatic capacity^1–3^, whether or how tumor evolution or the metastatic site itself influences their contribution to tumor maintenance is unknown. Such knowledge would have important implications for precision oncology and may guide the development of much needed metastasis-targeting therapies.

Among driver mutations, inactivation of the *SMAD4* tumor suppressor gene – a core mediator of TGFβ signaling – is a hallmark of several gastrointestinal malignancies that is found at highest frequency in PDAC^4,5,10^. During cancer progression, *SMAD4* inactivation shifts TGFβ’s activity from tumor-suppressive to tumor-promoting by impairing its ability to trigger cell cycle arrest and EMT-coupled apoptosis^9,23^. Accordingly, *SMAD4*-mutant tumors have higher rates of metastasis in PDAC patients, an effect recapitulated in animal models^7,8,11,14^. However, it is unknown whether *SMAD4* inactivation maintains disease at metastatic sites, which is key to understand given that most PDAC patients are diagnosed after tumor cells have spread to distant organs^24^. In this study, we took advantage of a new murine model that enables inducible and reversible *Smad4* inactivation at different PDAC stages to interrogate the ongoing need for *Smad4* inactivation in metastatic disease. Our results reveal a diametrically opposed role for *Smad4* inactivation in sustaining liver and lung metastases and establish a critical interplay between driver mutations and organ-specific chromatin states that contributes to the heterogeneity of cancers driven by identical genetic lesions.

## RESULTS

### *Smad4*-restorable genetically engineered mouse model of PDAC

To enable reversible *Smad4* inactivation in PDAC, we generated a genetically engineered mouse model (GEMM) that harbors pancreas-specific, single-copy, doxycycline (Dox)-inducible short hairpin RNA (shRNA) against *Smad4* (or against *Renilla* luciferase as a control) on the background of oncogenic *Kras^G12D^*(hereafter KC-shSmad4 and KC-shRen, respectively; see Methods) (**Fig. 1a**). This genetic strategy allows for tumor development in the setting of *Smad4* depletion (by Dox administration) and subsequent restoration of *Smad4* expression at physiological levels from its endogenous locus (by Dox withdrawal). The alleles also contain two fluorescent reporters that facilitate identification and isolation of tumor cells: a constitutive mKate2 and an shRNA-linked GFP (**Fig. 1a**).

**Figure 1.**
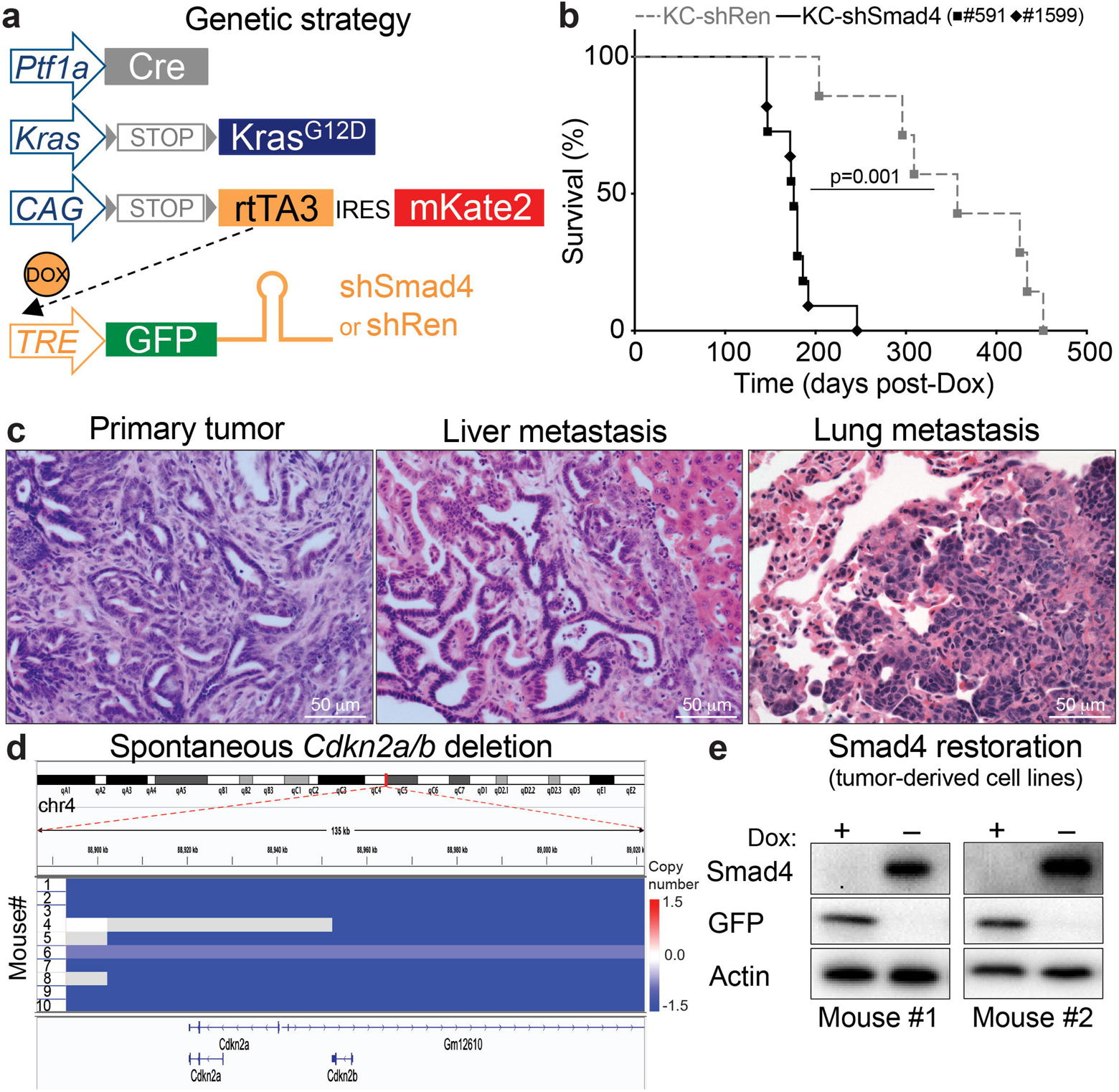
A *Smad4*-restorable genetically engineered mouse model of PDAC. **(a)** Schematic of GEMM alleles. rtTA3 = reverse tetracycline trans<u>a</u>ctivator, <u>3</u>^rd^ generation; Dox = <u>dox</u>ycycline; TRE = tetracycline response <u>e</u>lement. **(b)** Overall survival of KC-shRen mice and KC-shSmad4 mice (expressing one of two independent Smad4 shRNAs: #591 or #1599) after Dox administration (n=7 KC-shRen mice; n=11 KC-shSmad4 mice). Statistical analysis by log-rank test. **(c)** Representative hematoxylin and eosin (H&E) staining of primary tumors and liver and lung metastases from KC-shSmad4 mice. **(d)** Sparse whole-genome sequencing (sWGS) analysis of the *Cdkn2a/b* locus in KC-shSmad4 tumor-derived cell lines (n=10 independent mice). **(e)** Representative Western blot analysis of Dox response *in vitro* in two independent KC-shSmad4 cell lines.

Consistent with results from conventional knockout GEMMs^6,12–14^, *Smad4* depletion promoted tumor initiation, shortened survival, and led to the development of tumors that metastasize to the liver and, less frequently, to the lungs (**Fig. 1b, c; ED Fig. 1a**). At late stage, primary and metastatic tumors expressed the fluorescent reporters (**ED Fig. 1b**) and maintained potent depletion of SMAD4 protein (**ED Fig. 1c**). Notably, tumor formation appeared to require further inactivation of the *Cdkn2a* tumor suppressor gene, as sparse whole-genome sequencing of tumor-derived cell lines revealed spontaneous homozygous deletion of the *Cdkn2a/b* locus in 9/10 cases^25^ (**Fig. 1d**). This lesion (along with *Kras* gain) was the most prominent event in an otherwise largely stable genome (**ED Fig. 1d**), in agreement with the previously reported genome evolution of PDAC driven by alterations in the TGFβ pathway^26^. The *Cdkn2a/b* deletions were highly concordant between primary and metastatic tumors (**ED Fig. 1e**), and they mirrored the genetic association of *SMAD4* alterations with homozygous *CDKN2A/B* deletions in the MSK IMPACT cohort of human PDAC (**ED Fig. 1f**). Thus, the KC-shSmad4 GEMM recapitulates cardinal features of the human disease and further enables reversible *Smad4* inactivation.

### *Smad4* restoration has organ-specific effects on tumor growth

To study *Smad4* reactivation in metastatic tumors, we turned to transplantation assays using cell lines derived from primary GEMM tumors that were capable of metastasis, as this approach afforded longer experimental time before tumor burden necessitated mouse euthanasia. The tumor-derived cell lines maintained robust *Smad4* restorability (**Fig. 1e**), mounted an expected *Smad4*-dependent cytostatic response to TGFβ *in vitro* (**ED Fig. 2a, b**), and produced metastases with remarkably similar histopathology to those emerging in the GEMMs and in PDAC patients^27^ (**ED Fig. 2c**). *Smad4*-dependent apoptotic responses induced by TGFβ in PDAC epithelial progenitors^9,28^ were not captured in these cell lines at the analyzed timepoints.

KC-shSmad4 cells were stably transduced with firefly luciferase to facilitate tumor monitoring and delivered via orthotopic, intrasplenic, or tail vein injections into nude mice to respectively generate primary tumors, liver metastases, or lung metastases (**Fig. 2a**). After 4-6 weeks, which allowed for tumor formation under *Smad4*-depleted conditions (+Dox = Smad4 OFF), *Smad4* expression was restored by Dox withdrawal (–Dox = Smad4 ON) in a randomly selected half of each cohort, and tumor burden was assessed 30 days later (**Fig. 2a**). Strikingly, the response to *Smad4* restoration was different in each of the three organs: tumor burden was unchanged in the pancreas, decreased in the liver, and increased in the lungs (**Fig. 2b-d; ED Fig. 2d**). Of note, similar results were obtained in spontaneous metastases arising from the orthotopic KC-shSmad4 transplants but not in KC-shRen cells, thus ruling out artifactual effects due to the employed metastasis assays or Dox administration (**ED Fig. 2e-g**).

**Figure 2.**
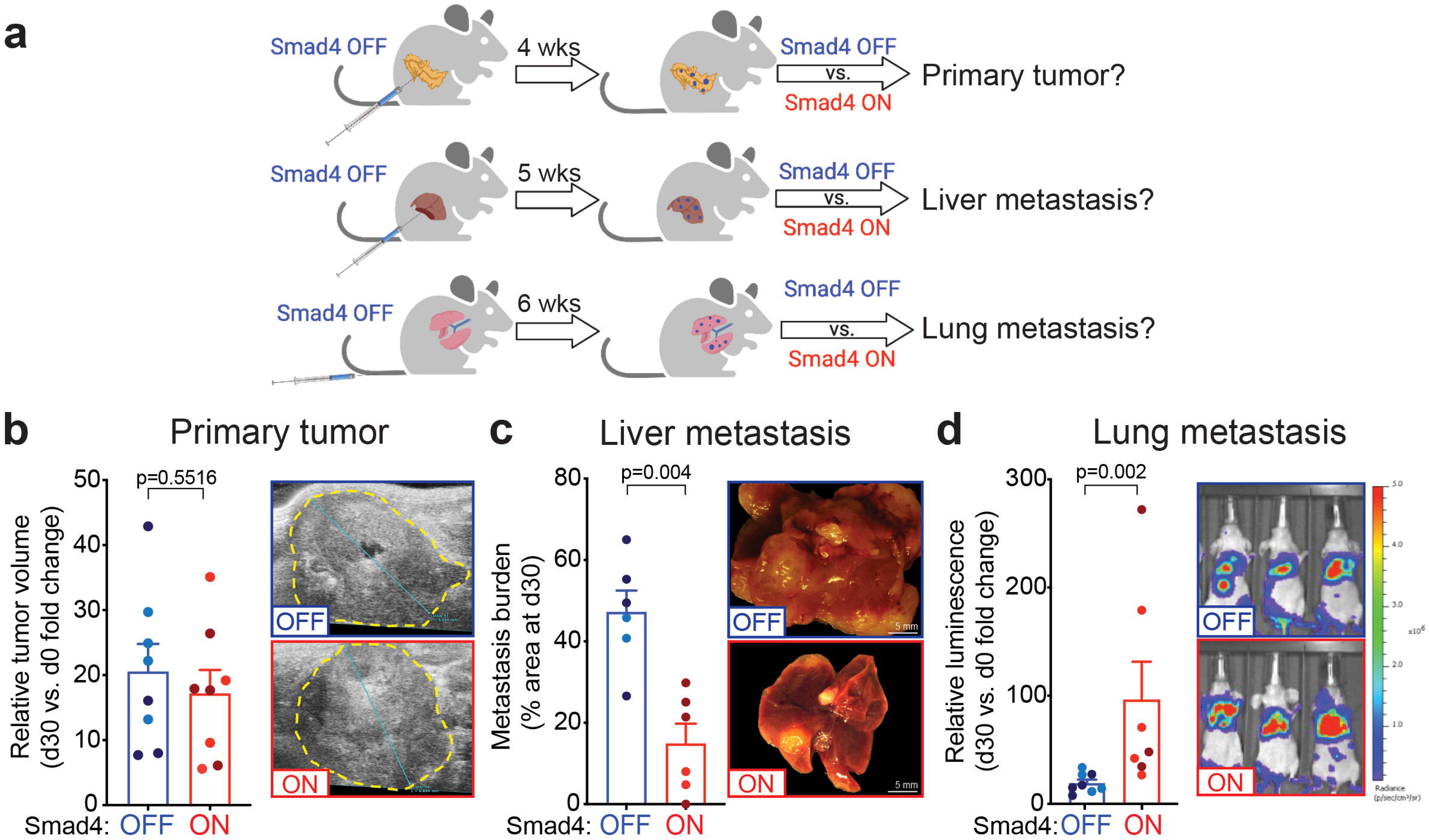
*Smad4* restoration has organ-specific effects on tumor growth. **(a)** Schematic of orthotopic, intrasplenic, and tail vein injection experiments. **(b)** Analysis of primary tumor growth after orthotopic transplantation of KC-shSmad4 cells and subsequent *Smad4* restoration. (Left) Fold-change quantifications of tumor volume at day 30 vs. day 0 of Dox withdrawal (n=8 independent mice per group). Different color shading indicates independent cell lines. (Right) Representative ultrasound images of tumors (demarcated by dashed yellow lines). **(c)** Analysis of liver metastasis burden after intrasplenic injections of KC-shSmad4 cells and subsequent *Smad4* restoration. (Left) Percent-area quantifications at day 30 after Dox withdrawal (n=6 independent mice per group). Different color shading indicates independent cell lines. (Right) Representative macroscopic images of tumor-bearing livers. **(d)** Analysis of lung metastasis burden after tail vein injections of KC-shSmad4 cells and subsequent *Smad4* restoration. (Left) Fold-change quantifications of bioluminescent signal at day 30 vs. day 0 of Dox withdrawal (n=8, 7 independent mice per group, respectively). Different color shading indicates independent cell lines. (Right) Representative bioluminescent images of tumor-bearing mice. Statistical analysis: (b) Unpaired two-tailed t-test; (c, d) Mann-Whitney test.

To determine the relevance of our findings to human PDAC, we queried data on metastatic recurrence in PDAC patients after primary tumor removal, where *SMAD4* status was evaluated by immunohistochemistry (IHC) staining of resected primary or metastatic tumors^29,30^. Corroborating a potentially tumor-suppressive vs. promoting function of *SMAD4* in the liver vs. lungs, 69% of cases with recurrent liver metastases lacked SMAD4 expression, in contrast to 50% of concurrent and 33% of isolated lung metastases (**ED Fig. 2h**). Thus, *Smad4* inactivation confers a selective advantage to liver metastases and a disadvantage to lung metastases, consistent with SMAD4 expression patterns in PDAC patients with metastatic recurrence.

### *Smad4* induces different transcriptional programs in liver vs. lung metastases

SMAD4 acts as a transcription factor (TF) by forming a complex with the SMAD2/3 TFs to activate gene expression programs downstream of TGFβ receptors^31^. Hence, we performed RNA sequencing (RNA-seq) to explore the transcriptional basis of the observed organ-specific phenotypes. Leveraging the mKate2 reporter in our model, we used fluorescent-activated cell sorting (FACS) to isolate tumor (mKate2^+^) cells 7 and 14 days after Dox withdrawal, which allowed for assessment of *Smad4* output kinetics. Consistent with its organ-specific effects on tumor growth, *Smad4* restoration led to upregulation of partially overlapping but mostly distinct genes, as compared to tumor cells kept on Dox: 89% (1580/1773) and 74% (613/825) organ-specific genes at days 7 and 14, respectively (**ED Fig. 3a**). This phenomenon was particularly pronounced in the liver at the 7-day timepoint and was still observed across all three organs at day 14 (**Fig. 3a**). Importantly, these results were not confounded by differential *Smad4* expression or baseline TGFβ signaling, since the extent of *Smad4* depletion/restoration and the levels of phosphorylated SMAD2 (a SMAD4-independent readout of TGFβ signaling^31^) were similar between the three organs (**ED Fig. 3b, c**).

**Figure 3.**
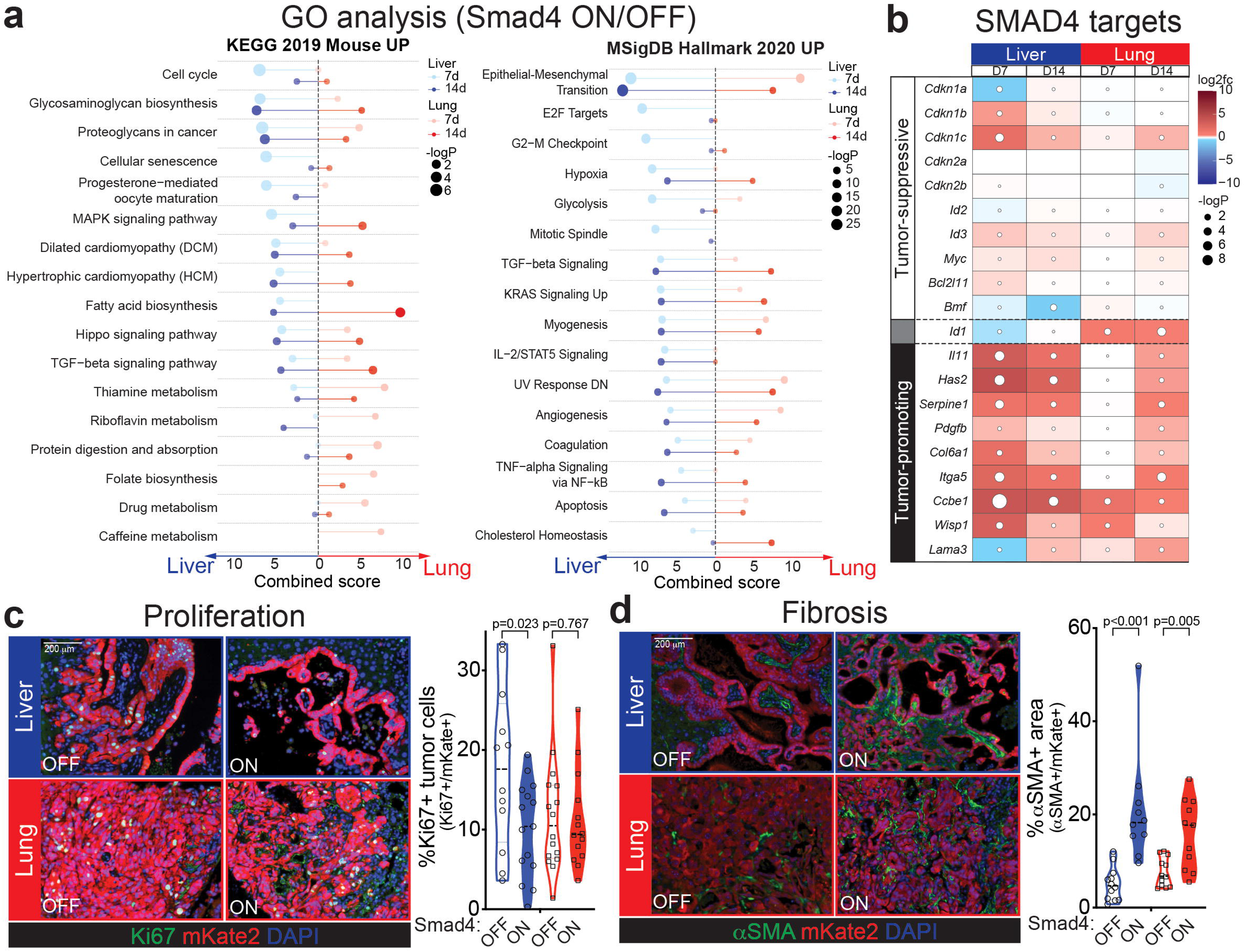
*Smad4* induces different transcriptional programs in liver vs. lung metastases. **(a)** Gene ontology (GO) analysis of upregulated genes in liver or lung metastases at 7 or 14 days after Dox withdrawal. Combined scores and p-values for the top KEGG (left) or Hallmark (right) pathways are shown for Smad4 ON vs. OFF in each organ and timepoint. Complete GO lists are provided in ED Table 1. **(b)** Heatmap of representative genes from *Smad4*’s cytostatic/apoptotic (tumor-suppressive) and fibrogenic (tumor-promoting) transcriptional programs. Average RNA-seq log_2_ fold-change and p-values are shown for Smad4 ON vs. OFF in each organ and timepoint **(c-d)** Representative immunofluorescence staining for Ki67 (c) or α-SMA (d) in KC-shSmad4 liver and lung metastases +/-*Smad4* restoration. mKate2 labels tumor cells. Quantifications are shown on the right (n=10-16 independent tumors from 3-4 mice). Ki67 analysis was performed 7 days after Dox withdrawal; α-SMA analysis was performed at experimental endpoint (30 days for liver; 45 days for lungs). Statistical analysis by unpaired two-tailed t-test (Ki67) or Mann-Whitney test (α-SMA).

To better understand the organ-specific consequences of *Smad4* restoration, we next performed functional annotation of *Smad4*-activated genes in the liver and lungs, as these organs exhibited opposite tumor growth phenotypes. Gene ontology analysis revealed that tumors in both organs upregulated extracellular matrix (e.g. glycosaminoglycan, proteoglycan, and focal adhesion) and EMT-related transcriptional programs, while only liver metastases showed enrichment for cell cycle and senescence-related gene signatures (**Fig. 3a; ED Fig. 3d**). Intersection of these gene lists with available data from SMAD2/3 ChIP-seq (chromatin immune-precipitation followed by sequencing) of murine PDAC cells^9,28^ confirmed differential engagement of SMAD4-dependent binding targets, further implicating direct effects of *Smad4* reactivation (**ED Fig. 3e, f**).

Importantly, the differentially expressed targets included genes that distinguish between SMAD4’s tumor-suppressive and tumor-promoting functions. They prominently featured upregulation of the tumor suppressor gene and cell cycle inhibitor *Cdkn1c* (also known as p57^KIP2^)^32,33^ specifically in the liver, as well as induction of SMAD4’s tumor-promoting fibrogenic program (including *Il11, Has2, Serpine1, Col6a1, Itga5, Ccbe1* and *Wisp1*)^28^ in both organs, albeit with a delayed kinetics in the lungs (**Fig. 3b**). Of note, the SMAD4-dependent target *Id1*, known to reflect TGFβ’s pro-tumor mode of action^34^, showed elevated expression in lung metastases but downregulation in liver metastases (**Fig. 3b**).

These transcriptomic data suggesting differential cytostatic and fibrogenic outputs were validated by immunostaining for the proliferation marker Ki67 and the TGFβ-dependent myofibroblast marker α-smooth muscle actin (α-SMA)^28^. In support of a liver-specific cytostatic response and liver/lung-shared fibrogenic response, *Smad4* reactivation reduced the proportion of Ki67^+^ tumor cells only in the liver, while both organs exhibited increases in α-SMA levels (**Fig. 3c, d**). Taken together, our analysis reveals discordant engagement of *Smad4*’s tumor-suppressive vs. promoting effectors in liver vs. lung metastases, thus demonstrating that anatomical site can uncouple TGFβ’s anti- and pro-tumorigenic programs.

### Liver and lung metastases harbor distinct chromatin states

Given the organ-specific transcriptional responses, we hypothesized that liver and lung metastases may harbor distinct chromatin states that afford different accessibility to SMAD4’s target genes. To test this, we performed ATAC-seq (assay for transposase-accessible chromatin using sequencing) on FACS-isolated tumor cells from the pancreas, liver, and lungs +/- *Smad4* restoration. Unsupervised clustering of differentially accessible peaks linked distinct chromatin states to tumors residing in different organs, while *Smad4* status itself had a limited impact (**Fig. 4a; ED Fig. 4a, b**), consistent with the fact that it is not a pioneer TF^31,35^. However, the differential chromatin accessibility affected SMAD4-dependent target genes, including its cytostatic and fibrogenic effectors highlighted earlier (**ED Fig. 4c**).

**Figure 4.**
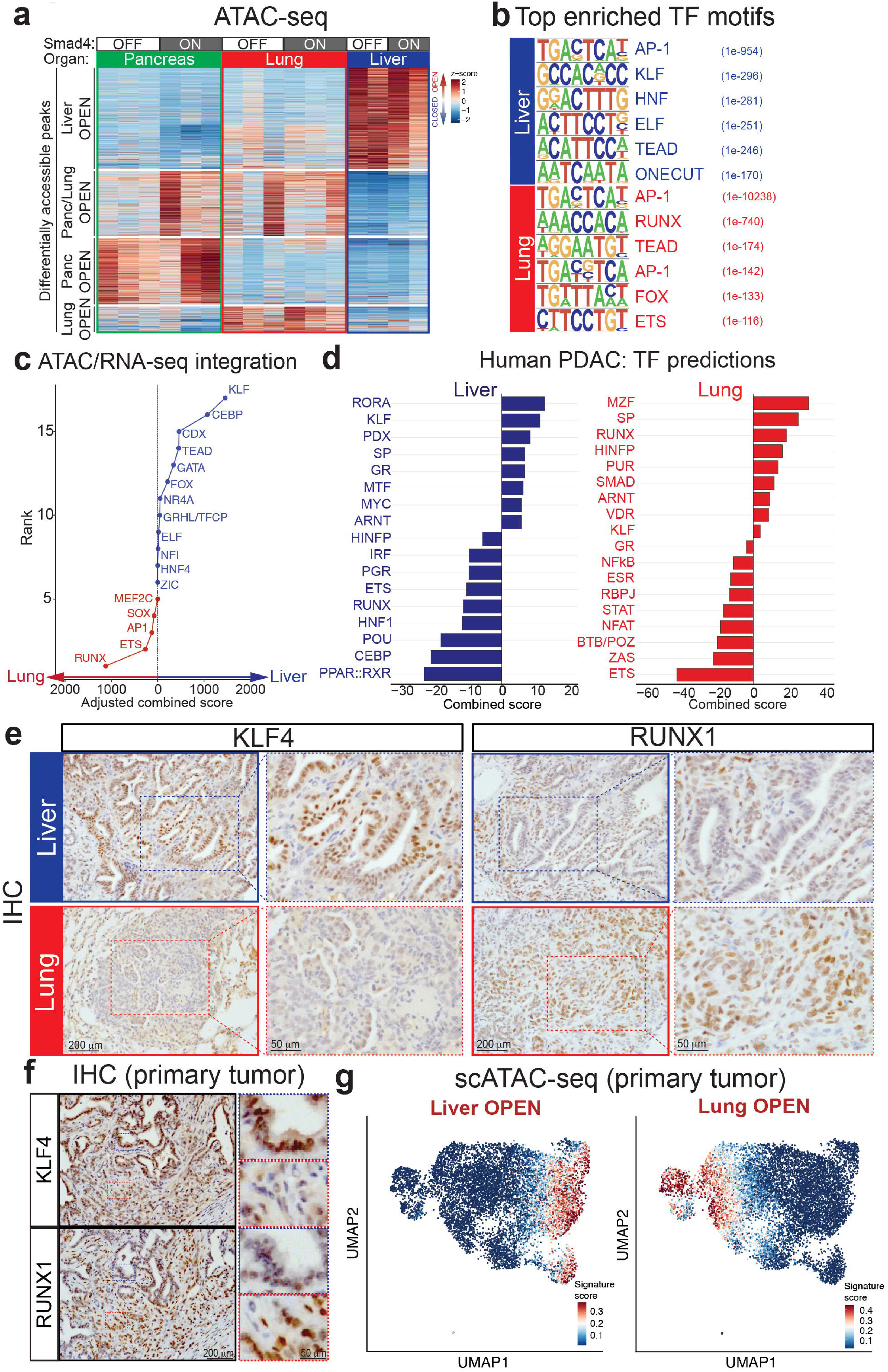
Liver and lung metastases harbor distinct chromatin states. **(a)** Heatmap of differentially accessible chromatin regions in tumor (mKate2^+^) cells from the pancreas, liver, or lungs. Smad4 ON corresponds to 14 days of Dox withdrawal. Each column represents an independent mouse. A complete list of differentially accessible ATAC-seq peaks is provided in ED Table 3. **(b)** Top-scoring TF motifs identified by HOMER *de novo* motif analysis of ATAC-seq peaks enriched in liver vs. lung metastases. The numbers in parentheses indicate enrichment p-values. **(c)** ATAC-RNA combined scores for the indicated TF families in liver vs. lung metastases. This metric infers the probability that a given TF family with a significantly enriched motif in the ATAC-open regions impacts SMAD4-induced gene expression changes, based on a consistent RNA-seq change in the Smad4 ON vs. OFF comparison (see Methods for details). Top TFs scored using the HOMER *de novo* motif analysis in liver vs. lungs are shown. **(d)** RNA-seq combined scores for the indicated TF families in liver and lung metastases from human PDAC patients^38^. This metric infers the activity of a given TF family, based on enrichment/depletion of its predicted target genes in the respective metastases vs. primary tumors but not in the corresponding normal tissues (see Methods for details). Top TFs scored using JASPAR/TRANSFAC position weight matrix (PWM) data are shown. **(e)** Representative IHC staining for KLF4 and RUNX1 in KC-shSmad4 (+Dox) liver or lung metastases. Higher magnifications of dashed areas are shown on the right to highlight tumor-specific nuclear signal. Data are representative of 20 metastases across 4 independent mice. **(f)** Representative IHC staining for KLF4 and RUNX1 in serial sections of a KC-shSmad4 (+Dox) primary tumor. Higher magnifications of dashed areas are shown on the right to highlight mutual exclusivity. Data are representative of 4 independent mice. **(g)** UMAP (uniform manifold approximation and projection) visualization of scATAC-seq profiles of KC-shSmad4 (+Dox) primary tumor cells (mKate2^+^). Signature scores based on liver- or lung-specific ATAC-open peaks from bulk ATAC-seq data are displayed in color per individual cell. Data are representative of 3 independent mice.

In light of these data, we then asked which other TFs are predicted to bind in the liver- vs. lung-specific ATAC-open regions and may thereby facilitate SMAD4’s organ-specific activity. Motif analysis revealed mostly distinct TF families, the top unique predictions being KLF, HNF, and ELF in the liver, and RUNX, FOX, and ETS in the lungs (**Fig. 4b**). To determine which of these may cooperate with SMAD4 to impact organ-specific gene expression, we integrated our RNA- and ATAC-seq datasets to identify TF families: (i) whose motifs were enriched in the differentially accessible regions, and (ii) whose predicted targets were upregulated upon *Smad4* restoration. This analysis narrowed down the list of organ-specific candidates to the KLF family in the liver and the RUNX family in the lungs (**Fig. 4c**), both of which are TF families with pioneer factor capabilities^35–37^. Finally, to assess which TF families are of highest relevance to human PDAC, we used RNA-seq data from metastatic human PDAC^38^ to impute differential TF activity in liver or lung metastases based on enrichment or depletion of a given TF’s target genes relative to primary tumors (see Methods for details). Corroborating our mouse data, KLF targets were enriched in liver metastases, while RUNX targets were depleted in liver and enriched in lung metastases (**Fig. 4d**). Overall, these results nominate the KLF and RUNX TF families as organ-specific determinants of chromatin-directed transcriptional programs responsive to SMAD4.

The KLF and RUNX families contain multiple TFs that have been implicated in PDAC biology. Among them, KLF4 and KLF5 are both enforcers of pancreatic epithelial identify^9,39–41^, whereby KLF5 silencing by TGFβ compromises PDAC cell survival^9^. Interestingly, KLF4 can exhibit pro- or anti-tumorigenic effects in PDAC depending on context, despite binding to similar DNA motifs as KLF5^9,41,42^. On the other hand, RUNX1 and RUNX3 have been implicated as drivers of invasion and metastasis and as potential genetic dependencies in PDAC^14,43,44^. To define which specific TF(s) are most likely to underlie the observed organ-specific phenotypes, we performed IHC staining for each of these factors in KC-shSmad4 metastases. KLF4 exhibited strong specificity for liver and RUNX1 for lung metastases; at the same time, KLF5, KLF6, and RUNX2 presented signal in both organs, albeit to a different extent, and RUNX3 did not show tumor-specific signal but rather stained stromal cells in our model (**Fig. 4e; ED Fig. 4d**). Thus, our refined analysis further nominates KLF4 and RUNX1 as particular TFs that are likely to mediate organ-specific chromatin opening in liver vs. lung metastases.

### Liver and lung metastasis-like cell states emerge in primary tumors

A growing body of evidence suggests that pro-metastatic epigenetic programs can arise early during tumorigenesis^45,46^. Interestingly, IHC staining for KLF4 and RUNX1 in KC-shSmad4 primary tumors was heterogeneous, indicating broad expression of both TFs yet in apparently non-overlapping subsets of cells (**Fig. 4f**). These data suggested that the KLF and RUNX-associated chromatin states – and their opposite responsiveness to SMAD4 – may already be present in sub-populations within the primary tumor. To further address this, we integrated our ATAC- and RNA-seq data from primary tumors +/- *Smad4* restoration (for 14 days) to infer KLF and RUNX TF activities. The latter were defined by combining metrics of chromatin accessibility at the respective TF motif with transcriptional changes in the TF’s target genes (see Methods for details). This analysis revealed that *Smad4* restoration caused a decrease of inferred KLF activity and an increase of inferred RUNX activity in the pancreas (**ED Fig. 4e**). These data support the concept that the KLF and RUNX-associated cell states pre-exist in the primary tumor, whereby SMAD4 antagonizes KLF activity while cooperating with RUNX activity.

To further assess whether chromatin states that pre-exist in the primary tumor are linked to organ-specific metastasis, we performed coupled single-cell multiomics (ATAC/RNA-seq) on three independent KC-shSmad4 primary tumors and then mapped liver- and lung-specific open chromatin peaks identified in our bulk ATAC-seq on the single-cell space. This analysis identified distinct primary tumor cell sub-populations that harbored chromatin states resembling the organ-specific states observed in established liver and lung metastases (**Fig. 4g; ED Fig. 4f**). We also probed previously published single-cell ATAC-seq data from pre-malignant pancreatic tissue (harboring only *Kras^G12D^*)^45^. Remarkably, this analysis yielded similar results to the advanced KC- shSmad4 primary tumors, implying that the organ-specific chromatin states may arise even before cells acquire malignant potential (**ED Fig. 4g**).

Finally, we asked if the identified liver- and lung-specific metastatic states can also be found in human PDAC. To this end, we leveraged the multiomic nature of our mouse single-cell data to generate matching RNA-seq signatures of the cell populations that were enriched for liver- vs. lung-specific chromatin peaks in the ATAC-seq analysis. The resulting signatures were then used to query single-cell RNA-seq data from 16 human primary PDAC samples^47^. This analysis confirmed the existence of distinct cell sub-populations in the human primary tumors that resemble the organ-specific cell states in our mouse model (**ED Fig. 4h**). The resemblance was particularly strong for the liver-specific states, likely because liver metastasis is more common than lung metastasis in PDAC patients. Overall, our data suggest that the liver and lungs favor metastatic cells harboring different chromatin states which are similar to pre-existing states found in the primary tumor.

### *Klf4* depletion is sufficient to reverse *Smad4* function in liver metastasis

Finally, we functionally interrogated the *Smad4*-TF interplay, focusing on the setting of liver metastasis where *Smad4*-mediated tumor suppression could be successfully restored. To do so, we used stable shRNA-based knockdown to deplete *Klf4* or *Runx1* (or shRNAs to target *Renilla* luciferase as a neutral control) in KC-shSmad4 cells. While both sh*Klf4* and sh*Runx1* achieved potent reduction of the respective proteins (**Fig. 5a**), the corresponding cell lines had unaltered basal proliferation and response to TGFβ upon *Smad4* restoration *in vitro* in comparison to the sh*Ren* control (**ED Fig. 5a**). We then generated liver metastases by intrasplenic injection of mice on Dox (Smad4 OFF), as described earlier (**ED Fig. 5b**). Consistent with a general role for *Klf4* in supporting a liver-metastatic cell state, its depletion reduced baseline tumor burden; by contrast, *Runx1* depletion produced a smaller reduction in tumor burden that did not reach statistical significance (**ED Fig. 5c**).

**Figure 5.**
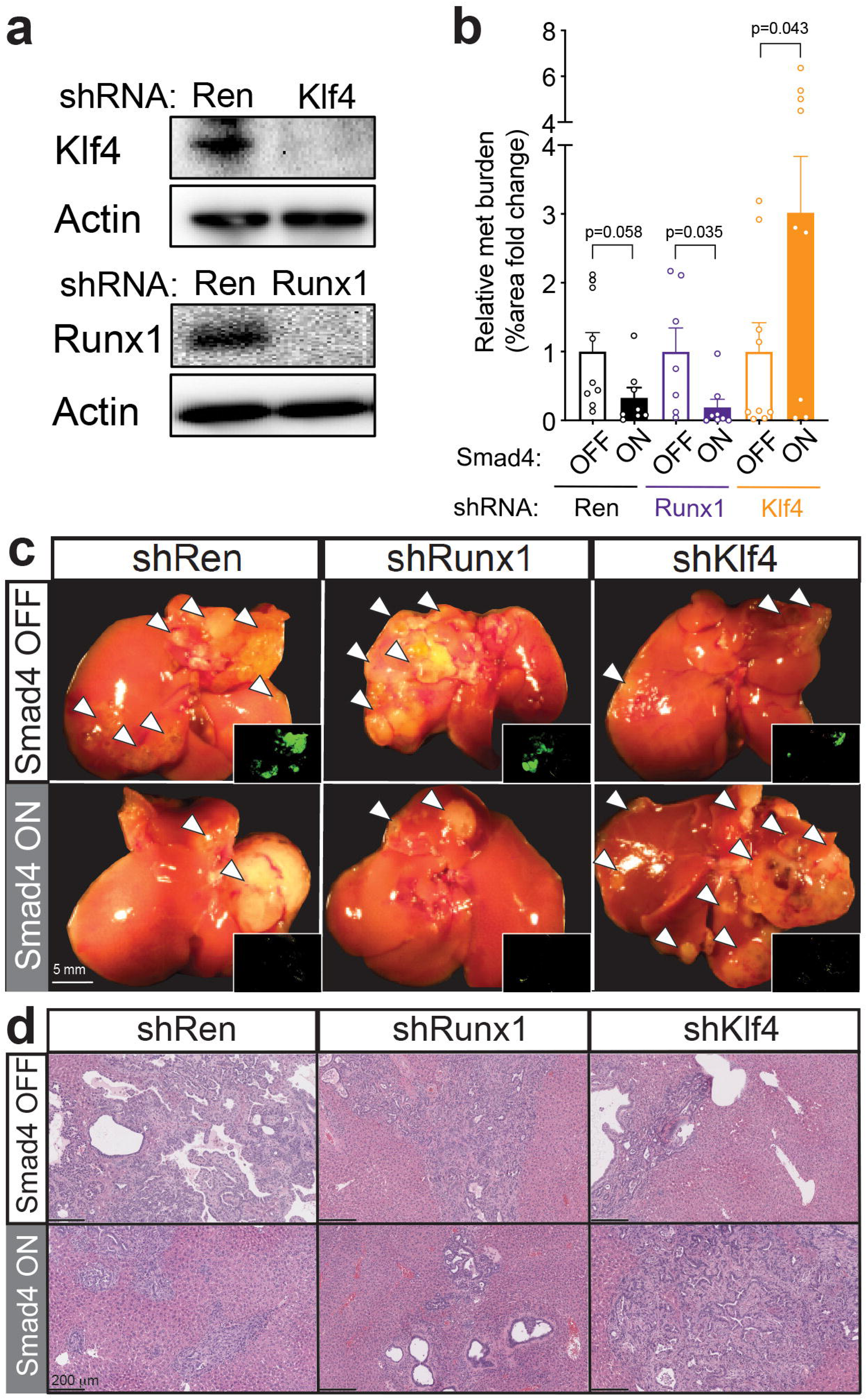
*Klf4* depletion is sufficient to reverse *Smad4* function in liver metastasis. **(a)** Western blot analysis of shRNA efficiency. Actin was used as a loading control. **(b)** Quantifications of metastasis burden (fold-change of % tumor area) at day 30 after Dox withdrawal (n=8, 8, 7, 8, 9, 9 independent mice per group, respectively). Statistical analysis by unpaired two-tailed t-test. **(c)** Representative macroscopic images of tumor-bearing livers +/-*Smad4* restoration on the background of the indicated shRNA-based knockdowns. Inset shows shSmad4-linked GFP reporter. Data are representative of the number of independent mice per group specified in (b). **(d)** Representative H&E staining of tumor-bearing livers +/-*Smad4* restoration on the background of the indicated shRNA-based knockdowns. Data are representative of 6 independent mice per group.

The effect of *Klf4* depletion on the tumor-suppressive response to *Smad4* in liver metastasis was striking. Whereas the sh*Ren-* and sh*Runx1*-expressing metastases displayed expectedly similar tumor suppression upon *Smad4* reactivation, sh*Klf4-*expressing metastases now exhibited a tumor-promotion phenotype of a three-fold increase in metastasis burden (**Fig. 5b-d**). Thus, *Klf4* depletion is sufficient to reverse *Smad4* function in liver metastases, which demonstrates that KLF4 facilitates *Smad4*’s tumor-suppressive activity in this setting. These data provide functional validation of our model whereby KLF4 cooperates with *Smad4* inactivation to sustain liver metastases (**ED Fig. 5d**).

## DISCUSSION

Our results demonstrate how driver gene mutations that are important for tumor initiation can show opposite requirements for maintenance at metastatic sites and place the paradigmatic duality of TGFβ signaling in an anatomical context. While it has been established that *SMAD4* mutation can switch TGFβ’s activity from tumor-suppressive to tumor-promoting^23^, here we show that this switch can also be mediated by the organ location of otherwise isogenic tumors. *SMAD4* loss itself thus does not universally favor tumor growth since its inactivation appears to be a liability rather than an advantage for lung metastases, at least in the clinically relevant setting of pre-seeded metastases modeled in our study. Hence, our data provide a possible reason for the unusually low rates of *SMAD4* inactivation in isolated lung metastases of PDAC patients^29,30,48^.

Mechanistically, the involvement of KLF and RUNX factors is consistent with both the operational logic of TGFβ signaling – whose contextual effects are often defined by interplay with such developmental TFs^31^ – and the reported functions of these TFs in PDAC and other settings. In particular, certain KLF factors promote the epithelial cell fate^39^, which in turn is known to be essential for colonization of the liver^49,50^, whereas RUNX proteins facilitate extracellular matrix remodeling that can create a pro-metastatic fibrogenic microenvironment in the lungs when unopposed by a tumor-suppressive program^46^. Future studies will determine the extent to which organ-specific effects apply to other common cancer drivers, many of which also enhance metastatic proclivity; such drivers include missense mutations of the *TP53* tumor suppressor gene^51^, gains/amplifications of the mutant-*KRAS/PTHLH*^26,52^ and *MYC*^53^ loci, or deletions of the *CDKN2A/*type I interferon locus^25^.

Our results also add to the growing appreciation that tissue context can influence the output of gene mutations in cancer, for example, as illustrated by the differential susceptibility of cells from particular tissues or tissue locations to certain oncogenic events^54–57^. Our findings extend this concept to metastasis by showing how organ site can have a profound impact on a single driver mutation in a tumor from the same tissue of origin. As an underlying mechanism, we show that such mutations synergize with or antagonize distinct chromatin states that emerge early during tumorigenesis and are favored by different metastatic sites. Additional work will define the contribution of immune and other factors in the organ microenvironment that likely influence this gene–chromatin interplay. Regardless, when extended to precision oncology, our results draw attention to potentially divergent responses to therapy based on the tumor’s organ of residence. As such, they are in line with clinical observations of organ heterogeneity in therapy response^58–61^ and invite consideration of organ-specific therapies for tumors driven by mutations that show such dependence on metastatic site.

## METHODS

### Animals and *in vivo* procedures

#### Animal care

All mouse experiments were approved by the Memorial Sloan Kettering Cancer Center Institutional Animal Care and Use Committee (IACUC). Mice were maintained under pathogen-free conditions, housed on a 12 h–12 h light–dark cycle under standard temperature and humidity of approximately 18–24°C and 40–60%, respectively. Food and water were provided *ad libitum*. GEMMs were generated in house. *Foxn1^nu^* (athymic nude) mice used for transplants were purchased from Envigo or The Jackson Laboratory.

#### GEMMs

*Ptf1a^Cre/+^_;_LSL-Kras^G12D/+^_;_Rosa26^LSL-rtTA3-IRES-mKate2/+^(RIK);Col1a1^shRNA-Homing-Cassette/+^(CHC)* embryonic stem cells (ESCs)^62^ were targeted with two independent GFP-linked Smad4 shRNAs (shSmad4.591: CAAAGATGAATTGGATTCTTT; shSmad4.1599: ACAGTTGGAATGTAAAGGTGA) cloned into miR30-based targeting constructs, as previously described^62,63^. Targeted ESCs were selected and functionally tested for single integration of the GFP-linked shRNA element into the CHC locus, as previously described^62^. The KC-shRen ESC control clone used in this study has been described^62^. Before injection, ESCs were expanded briefly in KOSR+2i medium^64^ and confirmed to be negative for mycoplasma. Mice were generated by 8-cell or blastocyst injection of KC-shSmad4 or KC-shRen ESCs, and short hairpin RNAs were induced by treatment of the resulting mice with doxycycline (625 mg/kg) in the drinking water starting at 5-6 weeks of age. The identity of the ESCs and ESC-derived mice were authenticated by genomic PCR using a common Col1a1 primer paired with an shRNA-specific primer, all yielding products of approximately 250 bp:

- Col1a1: 5’-CACCCTGAAAACTTTGCCCC-3’;
- shRen.713: 5’-GTATAGATAAGCATTATAATTCCTA-3’;
- shSmad4.591: 5’- GTATAAAGAATCCAATTCATCTT-3’;
- shSmad4.1599: 5’-TATTCACCTTTACATTCCAAC-3’.

#### Orthotopic transplantation assays

Mouse hosts were placed on doxycycline chow 5-7 days before transplantation. Mice were anesthetized and a survival surgery was performed to expose the pancreas. 1 x 10^5^ tumor-derived cells were resuspended in 25 μL 1:1 DMEM (Gibco) : Matrigel (Corning) mix and injected in the tail of the pancreas of each mouse. Tumor engraftment and progression were monitored by palpation and ultrasound imaging. Where applicable, doxycycline withdrawal was done 4 weeks post-injection (corresponding to 3-5 mm tumor diameter) by switching the food source to regular chow. Primary tumor size was measured using ultrasound imaging (see below). At experimental endpoint, primary tumors, livers, and lungs were dissected and imaged under a dissection microscope for brightfield, mKate, and GFP fluorescence (Nikon SMZ1500 with NIS-Element software). Endpoint liver and lung metastasis burden were measured by calculating percent tumor area (mKate+) as a fraction of overall organ area. Euthanasia was performed upon reaching experimental or humane endpoints according to IACUC guidelines. All mice used were 6–8-week-old *Foxn1^nu^* females.

#### Experimental metastasis assays

For liver metastasis assays, mice were anesthetized and a survival surgery was performed to expose the spleen. 4 x 10^5^ tumor-derived cells resuspended in 20 μL PBS were injected in the splenic parenchyma of each mouse, followed by removal of the spleen and cauterization (splenectomy). Tumor engraftment and progression were monitored by palpation and ultrasound imaging. At experimental endpoint, livers were dissected and imaged under a dissection microscope for brightfield, mKate, and GFP fluorescence (Nikon SMZ1500 with NIS-Element software). Endpoint liver metastasis burden was measured by calculating percent tumor area (mKate+) as a fraction of overall organ area. For lung metastasis assays, mice were restrained, and 2.5 x 10^5^ tumor-derived cells resuspended in 250 μL PBS were injected in the tail vein of each mouse. Bioluminescent imaging was used to monitor tumor engraftment and progression, and to measure tumor burden (see below). Where applicable, doxycycline withdrawal was done at 5 weeks (intrasplenic) or 6 weeks (tail vein) post-injection by switching the food source to regular chow. The occasional mice that developed tumors outside of the respective target organs were excluded from the analysis. Euthanasia was performed upon reaching experimental or humane endpoints according to IACUC guidelines. All mice used were 6–8-week-old *Foxn1^nu^* females.

#### Animal imaging

For ultrasound, mice were anesthetized, then high-contrast ultrasound imaging was performed on a Vevo 2100 System with a MS250 13-to 24-MHz scanhead (VisualSonics). Images were acquired and tumor volume was measured using the Vevo 2100 software (VisualSonics). For bioluminescence, mice were injected with luciferin (5 mg/mouse, i.p.; Gold Technologies), anesthetized for 10 min, and then imaged on a IVIS Spectrum imager (Perkin Elmer). Images were acquired and bioluminescence signal was measured using the Living Image software (Perkin Elmer).

### Histological, immunohistochemistry (IHC), and immunofluorescence (IF) analyses

Tissues were fixed overnight in 10% neutral buffered formalin (Fisher Scientific #22-050-105), embedded in paraffin, and cut into 5-μm sections. Hematoxylin & eosin staining was performed using standard protocols. For immunostaining, slides were heated for 30 min at 55°C, deparaffinized, rehydrated with an alcohol series, and subjected to antigen retrieval with citrate buffer (Vector Laboratories #H-3300) for 25 min in a pressure cooker set on high. Sections were treated with 3% H_2_O_2_ for 10 min followed by a wash in deionized water (for IHC only), washed in PBS, then blocked in PBS/0.1% Triton X-100/1% BSA. Primary antibodies were incubated overnight at 4°C in blocking buffer. The following primary antibodies were used: mKate2 (Evrogen #AB233, 1:1,000, IF), SMAD4 (Millipore #04-1033, 1:200, IHC), pSMAD2 (Cell Signaling #3108, 1:100, IHC), Ki67 (BD Pharmingen #550609, 1:200, IF), α−SMA (Sigma #A2547, 1:1,000, IF), KLF4 (Abcepta #AM2725A, 1:100, IHC), KLF5 (Abcam #ab137676, 1:500, IHC), KLF6 (Abcam #ab241385, 1:1,000, IHC), RUNX1 (Cell Signaling #8529, 1:500, IHC), RUNX2 (Cell Signaling #12556, 1:500, IHC), RUNX3 (Life Technologies #MA5-17169, 1:500, IHC).

For IHC, HRP-conjugated secondary antibodies (ImmPRESS kits, Vector Laboratories #MP7401 and #MP2400) were applied for 30-60 minutes at room temperature and visualized with DAB substrate (ImmPACT kit, Vector Laboratories #SK-4105). Tissues were then counterstained with hematoxylin, dehydrated, and mounted with Permount (Fisher Scientific #SP15-100). For IF, secondary Alexa Fluor 488 or 594 dye-conjugated antibodies (Life Technologies, 1:500) were applied for 60 min at room temperature. Tissues were then counterstained with DAPI and mounted with Prolong Gold Antifade Mountant (Life Technologies #P36930).

Images were acquired on a Zeiss AxioImager microscope using a 10X, 20X, or 40X objective, an ORCA/ER CCD camera (Hamamatsu Photonics), and ZEN 3.3 software (Zeiss). For pSMAD2 quantification, the number of pSMAD2+ cells per randomly chosen IHC-stained 40X field of view in a tumor region was manually counted. For Ki67 and α−SMA quantification, 20X fields of view co-stained for mKate2 were analyzed as follows: mKate2+ areas were selected and then (1) the corresponding Ki67+ cells were counted and calculated as a percentage of DAPI+ cells; or (2) the corresponding α−SMA+ area was measured and calculated as a percentage of the mKate2+ area. Image analyses were performed using ImageJ/FIJI (NIH, USA). The number of analyzed samples and statistical analyses used for each assay are specified in the respective figure legends.

### Cloning

ESC-targeting plasmids were generated as described under ‘GEMMs’ above. The firefly luciferase reporter plasmid (pMSCV-Luc2-Blast) was generated by subcloning Luc2 from the pCDH-EF1-Luc2-P2A-tdTomato plasmid into the pMSCV-Blasticidin retroviral vector at the BglII/HpaI restriction sites with the inclusion of a Kozak sequence (GCCACC) upstream of the ATG start codon, using standard protocols. Constitutive *Renilla*, *Klf4*, and *Runx1* shRNAs were cloned in the pMSCV-mirE-SV40-Neomycin-BFP retroviral vector^65^ at the XhoI/EcoRI restriction sites, using standard protocols. pCDH-EF1-Luc2-P2A-tdTomato was a gift from Kazuhiro Oka (Addgene #72486). pMSCV-Blasticidin was a gift from David Mu (Addgene #75085)^66^. All plasmids were authenticated by test digestion and Sanger sequencing. The following shRNA sequences were used:

- Ren: 5’-GTATAGATAAGCATTATAATTCCTA-3’
- Klf4: 5’-TATAAAAATAGACAATCAGCA-3’
- Runx1: 5’-AAATCAGAAGCATTCACAGTT-3’

### Cell culture

All cells were maintained in a humidified incubator at 37°C with 5% CO_2_.

#### Primary cell line derivation

Cell lines were generated from tumor-bearing pancreata, livers, or lungs of KC-shSmad4 or KC-shRen mice. Liver- and lung-derived lines were only used for sWGS analysis. Tumors were dissected, chopped with razor blades, and digested with 1 mg/ml collagenase V (Sigma-Aldrich) diluted in HBSS for 30-60 min, followed by 0.25% trypsin for 5-10 min. Digested tissues were washed with complete DMEM (DMEM, 10% FBS (Gibco), 1X penicillin–streptomycin), passed through a 100-μm filter, and cultured in complete DMEM on collagen-coated plates (PurCol, Advanced Biomatrix, 0.1 mg/ml) supplemented with 1 μg/ml doxycycline at 37°C. Cells were passaged at least five times to eliminate any non-tumor cells before using them in experiments. Primary cultures were authenticated by flow cytometry of engineered fluorescent alleles. All cultures were tested negative for mycoplasma.

#### Virus generation and transduction

For stable transduction of firefly luciferase and constitutive shRNA constructs, VSV-G pseudotyped retroviral supernatants were generated from transduced Phoenix-GP packaging cells and infections were performed as described elsewhere^67^. Infected cells were selected with 10 μg/ml Blasticidin for 5 days or 800 μg/ml G418 for 7 days, depending on the selection marker. Luciferase expression was confirmed by *in vitro* and *in vivo* bioluminescence. Knockdowns were confirmed by Western blot using standard procedures and the following antibodies: SMAD4 (Santa Cruz #sc-7966, 1:500), KLF4 (Abcepta #AM2725A, 1:1000), RUNX1 (Cell Signaling #8529, 1:1000), and Actin-HRP (Sigma #A3854, 1:20,000).

### Tumor cell isolation

For RNA-seq, ATAC-seq, and scMultiomics analyses, tumor cells were freshly isolated from pancreata, livers, or lungs of KC-shSmad4 mice by FACS. Specifically, dissected tumors were finely chopped with scissors and incubated with digestion buffer containing 1 mg/ml collagenase V (Sigma #C9263), 2 U/ml dispase (Life Technologies #17105041) dissolved in HBSS with Mg^2+^ and Ca^2+^ (Thermo Fisher Scientific #14025076) supplemented with 0.1 mg/ml DNase I (Sigma #DN25-100MG) and 0.1 mg/ml soybean trypsin inhibitor (STI) (Sigma #T9003), in gentleMACS C Tubes (Miltenyi Biotec) for 42 min at 37 °C using the gentleMACS Octo Dissociator. After enzymatic dissociation, samples were washed with PBS and further digested with a 0.05% solution of Trypsin-EDTA (Gibco #15400) diluted in PBS for 5 min at 37°C. Trypsin digestion was neutralized with FACS buffer (10 mM EGTA and 2% FBS in PBS) containing DNase I and STI. Samples were then treated with RBC Lysis Buffer (Invitrogen #00-4333-57) for 5 min at room temperature, washed in FACS buffer containing DNase I and STI, and filtered through a 100-μm strainer. Cells were then resuspended in FACS buffer containing DNase I, STI and 300 nM DAPI as a live-cell marker, and filtered through a 40-μm strainer. Cells were sorted on a BD FACSAria I, BD FACSAria III (Becton Dickinson), or MA900 (Sony) cell sorter for mKate2 (co-expressing GFP for on-Dox shRNA mice), excluding DAPI+ cells. Cells were sorted directly into TRIzol LS (Thermo Fisher Scientific # 10296028) for RNA-seq or collected in 2% FBS in PBS for ATAC-seq.

### Sparse whole-genome sequencing

Low-pass whole-genome sequencing was performed on genomic DNA freshly isolated from cultured cells as previously described^68^. Briefly, 1 μg of gDNA was sonicated on an E220 sonicator (Covaris; settings: 17Q, 75s), and libraries were prepared by standard procedure (end repair, addition of polyA, and adaptor ligation). Libraries were then purified (AMPure XP magnetic beads, Beckman Coulter), PCR enriched, and sequenced (Illumina HiSeq). Reads were mapped to the mouse genome, duplicates removed, and an average of 2.5 million reads were used for CNA determination with the Varbin algorithm^69^.

### RNA-sequencing

#### RNA extraction, RNA-seq library preparation and sequencing

Total RNA was isolated using TRIzol LS (Thermo Fisher Scientific # 10296028) followed by column clean-up using an RNeasy kit (Qiagen #74106). RNA was quantified using Nanodrop and its quality assessed by an Agilent 2100 BioAnalyzer. 100-500 ng of total RNA underwent polyA selection and TruSeq library preparation according to instructions provided by Illumina (TruSeq Stranded mRNA HT Kit, Illumina #20020595), with 15 cycles of PCR. Samples were barcoded and run on a HiSeq (Illumina) in a 75-bp SE run, with an average of 50 million reads per sample.

#### RNA-seq read mapping, differential expression analysis and heatmap visualization

RNA-seq data were analyzed by removing adaptor sequences using Trimmomatic^70^. RNA-seq reads were then aligned to GRCm38.91 (mm10) with STAR^63^, and transcript count was quantified using featureCounts^71^ to generate raw count matrix. Differential gene expression analysis was performed using the DESeq2 package^72^ implemented in R (http://cran.r-project.org/). Principal component analysis (PCA) was performed using the DESeq2 package in R. Differentially expressed genes (DEGs) were determined by >2-fold change in gene expression with Benjamini-Hochberg adjusted p-value <0.05. For heatmap visualization of DEGs, samples were z-score normalized and plotted using the ‘pheatmap’ package in R.

#### Functional annotations of gene sets

Pathway enrichment analysis was performed using the indicated gene sets in enrichR^73^. Significance of the tests was assessed using combined score, described as c = log(p) * z, where c is the combined score, p is Fisher’s exact test p-value, and z is z-score for deviation from expected rank (shown in Figure 3 and ED Table 1), as well as by adjusted p-values defined by enrichR.

#### Intersection of RNA-seq and publicly available ChIP-seq data

To analyze transcriptional dynamics of SMAD2/3/4-binding targets, we used two publicly available PDAC datasets^9,28^. First, we extracted SMAD4-dependent SMAD2/3 ChIP-seq peaks that were significantly enriched in both studies (based on p-value < 1e-8 and an enrichment cut-off of > 8-fold increase of SMAD2/3 ChIP signal). These peaks were then associated with genes based on UCSC.mm10.knownGene using ChIPseeker package^74^: they were analyzed for genic location (annotatePeaks) and the nearest TSS was identified in order to annotate the peak to that gene. Finally, the resulting gene list was intersected with SMAD4-depednent differentially expressed genes in the present study. Log2FoldChange values between Smad4 ON vs. OFF in each organ were plotted, and the type of genomic binding region was color-annotated on the left (ED Fig. 3e).

### Bulk ATAC-sequencing

#### Cell preparation, transposition reaction, ATAC-seq library construction and sequencing

A total of 60,000 mKate2^+^ cells were isolated by FACS, washed once with 50 μl cold PBS and resuspended in 50 μl cold lysis buffer^75^. Cells were then centrifuged immediately for 10 min at 500*g* at 4°C, and the pellet of nuclei was subjected to transposition with Nextera Tn5 transposase (Illumina #FC-121–1030) for 30 min at 37°C, according to the manufacturer’s instructions. DNA was eluted using a DNA Clean & Concentrator Kit in 21 μl elution buffer (Zymo Research #D4013). ATAC-seq libraries were prepared using the NEBNext High-Fidelity 2X PCR Master Mix (NEB M0541) as previously described^76^. Purified libraries were assessed using a Bioanalyzer High-Sensitivity DNA Analysis kit (Agilent). Approximately 50 million paired-end 150-bp reads (25 million each side) were sequenced per replicate on a HiSeq instrument (Illumina).

#### Mapping, peak calling and dynamic peak calling

FASTQ files were trimmed with trimGalore and cutadapt^77^, and the filtered, pair-ended reads were aligned to mm10 with Bowtie2^78^. Peaks were called over input using MACS2^79^, and only peaks with a p-value of ≤ 0.001 and outside the ENCODE blacklist region were kept. All peaks from all samples were merged by combining peaks within 500 bp of each. featureCounts^71^ was used to count the mapped reads for each sample. The resulting peak atlas was normalized using DESeq2^72^. For comparison to DepthNorm, samples were normalized to 10 million mapped reads. Normalized bigWig files were created using the normalization factors from DESeq2 as previously described^80^ and BEDTools genomeCoverageBed^81^. Dynamic ATAC peaks were called if they had an absolute log_2_-transformed fold change ≥ 0.58 and FDR ≤ 0.1.

#### ATAC-seq heat map clustering

The dynamic peaks determined by comparing pancreas, liver and lung +/-Dox (=Smad4 OFF/ON) conditions were clustered using *z*-scores and *k*-means of 4 and plotted using ComplexHeatmap^82^.

#### TF motif enrichment analyses

Motif enrichment analysis was performed on differentially expressed ATAC peaks between liver and lung with the HOMER *de novo* motif discovery tool^83^ using findMotifsGenome command with the parameters size = given and length = 8-12. Motif enrichment scores of the *de novo* predicted motifs identified from this analysis were calculated for ATAC gain or loss regions between liver and lung +/-Dox (=Smad4 OFF/ON) by applying the findMotifsGenome command with size = given and length = 8-12 in each peak-set.

#### Integration of ATAC-seq and RNA-seq data

We adapted a previously described workflow^84^ that combines RNA-seq and ATAC-seq data with TF motif information to predict dominant TFs in liver vs. lung, as follows. For each organ, differential gene expression analysis between DoxOFF and DoxON was first performed individually (pancreas, liver, or lungs). Next, the resulting gene lists were intersected to identify organ-specific Smad4-responsive targets. EnrichR^73^ was used to calculate enrichment scores for annotated TF targets using the ChEA_2016 database. RNA-score was defined as -log10(adjusted p-value). Separately, HOMER was used to compare ATAC-seq data between Liver(DoxON) and Lung(DoxON) samples to identify differentially expressed peaks (DEPs). Custom TF motifs were curated by combining all the pairwise DoxON comparisons between any two organs, and these motifs were then used to reannotate the DEPs to get consistent enrichment scores across known TFs. ATAC-score was defined as -log10(p-value). Finally, Combined-scores were calculated by multiplying the respective RNA-and ATAC-scores. To determine the net change in ATAC-RNA combined scores for KLF and RUNX in the pancreas (ED Fig. 4e), each TF’s RNA- and ATAC-scores were calculated for Pancreas(DoxON) and Pancreas(DoxOFF) samples, multiplied to obtain combined scores for each Dox condition, and then the DoxON combined score was subtracted from the DoxOFF combined score.

### scMultiome-sequencing

#### Cell preparation, transposition reaction, scATAC-seq library construction and sequencing

Single Cell Multiome ATAC + Gene Expression was performed with 10X Genomics system using Chromium Next GEM Single Cell Multiome Reagent Kit A (catalog #1000282) and ATAC Kit A (catalog #1000280) following Chromium Next GEM Single Cell Multiome ATAC + Gene Expression Reagent Kits User Guide and demonstrated protocol, Nuclei Isolation for Single Cell Multiome ATAC + Gene Expression Sequencing. Briefly, cells (viability 95%) were lysed for 4 min and resuspended in Diluted Nuclei Buffer (10X Genomics #PN-2000207). Lysis efficiency and nuclei concentration was evaluated on Countess II automatic cell counter by trypan blue and DAPI staining. 11,000 nuclei were loaded per transposition reaction, targeting recovery of 7,000 nuclei after sequencing. After transposition reaction, nuclei were encapsulated and barcoded. Next-generation sequencing libraries were constructed following User Guide and sequenced on an Illumina NovaSeq 6000 system.

#### Quality control and cell filtering

Nucleosome signal score and TSS enrichment score for each cell were computed. Criteria for retaining individual cells were: TSS enrichment score >0.3, a nucleosome signal score <1.5, total ATAC-seq counts between 1,000 and 200,000 (based on the 10X Cell Ranger ATAC-seq count matrix) and total RNA counts between 1,000 and 50,000.

#### scRNA-seq data preprocessing and cell annotation

Gene expression UMI count data were normalized using SCTransform, percent mitochondrial RNA content was regressed out, and PCA was performed on the SCTransform Pearson residual matrix using the RunPCA function in Seurat^86^. Nearest neighbors were identified using FindNeighbors with dims = 1:30. The R package BBKNN was used to remove batch effects between mouse samples and cell types were annotated using R packages celldex, SingleR, Azimuth, and custom gene sets^85,86^. Only cells annotated as Ductal or Acinar cells were retained for downstream analysis.

#### scATAC-seq data preprocessing

scATAC-seq peaks were identified using MACS2 (ref. 79) using default parameters. Peak calling was performed using the CallPeaks function in Signac. Any peaks overlapping annotated genomic blacklist regions for the mm10 genome were removed. Counts for the resulting peak set were quantified for each cell using the FeatureMatrix function in Signac. For scATAC-seq analysis, data were normalized using the RunTFIDF function, and the top features were identified using FindTopFeatures with min.cutoff = ’10’. The RunSVD function was used to create the LSI space, and the resulting visualization was generated with UMAP using dims = 2:30. Clusters were identified using the “algorithm = 3” option.

#### Mapping of bulk ATAC-seq signatures

Differentially accessible peaks from bulk ATAC-seq were used to overlap with accessible peaks from scATAC-seq using intersectbed from bedtools^81^. Peaks with at least 1-bp overlaps were kept, and the top 5000 scATAC-seq peaks sorted based on significance were used to calculate ATAC signature scores using AddChromatinModule from Signac. LiverOPEN and LungOPEN cells were identified based on the signature score, and mutual exclusivity was calculated using the Chi-squared test.

#### Differential gene expression of scRNA-seq data

FindMarkers function from Seurat^86^ with “min.pct = 0.1” was used to identify differentially expressed genes (DEGs) between LiverOPEN and LungOPEN cells identified from scATAC-seq data. These DEGs were used for calculating gene signature scores for human PDAC scRNA-seq data.

### Public scATAC-seq analysis

Raw scATAC-seq data from ref. 45 were downloaded from the Gene Expression Omnibus (GSE137069). We kept cells using filters “min.cells > 10, min.features < 200”. Peaks that overlap with annotated genomic blacklist regions for the mm10 genome were identified. We retained cells using filters “peak_region_fragments > 3000, peak_region_fragments < 20000, pct_reads_in_peaks > 15, blacklist_ratio < 0.05, nucleosome_signal < 4, TSS.enrichment > 2”. Data were normalized using the RunTFIDF function, and the top features were identified using FindTopFeatures. We utilized the RunSVD function to create the LSI space, and the resulting visualization was generated with UMAP using dims = 2:30. Clusters were identified using the following options: “algorithm = 3 and resolution = 0.5 ”. GeneActivity from Signac was used to derive an approximate gene activity matrix, and custom gene signatures were used to annotate cell types. Ductal and acinar cells were retained for downstream analysis. Differentially accessible peaks from bulk ATAC-seq were used to intersect with accessible peaks from scATAC-seq using intersectbed from bedtools^81^. Peaks with at least 1-bp overlap were kept, and top5000 scATAC-seq peaks sorted based on significance were used to calculate ATAC signature scores using AddChromatinModule from Signac.

### Human PDAC analysis

#### Metastasis recurrence analysis

Previously reported data^29,30^ were re-analyzed by site of recurrence (liver or lungs) and annotation of SMAD4 IHC status (positive or negative).

#### MSK-IMPACT analysis

Human datasets were obtained through the MSK Clinical Sequencing Cohort (MSK-IMPACT) via cBioPortal^87,88^. Samples were selected as follows: (1) Cancer Type: Pancreatic cancer, (2) Cancer Type Detailed: Pancreatic Ductal Adenocarcinoma, and (3) Genotype (SMAD4: MUT HOMDEL). Comparison of genomic annotations between SMAD4-altered and unaltered (i.e. the rest) samples, along with corresponding statistical analyses, were generated and visualized by the cBioPortal.

#### Bulk RNA-seq analysis

Bulk RNA-seq data from ref. 38 was retrieved from the Gene Expression Omnibus (GSE71729). First, we compared liver or lung metastasis samples to primary tumor samples to derive liver or lung metastasis-specific differentially expressed genes. Second, we performed analogous comparison of normal liver or lung vs. normal pancreas samples. The tumor and normal gene sets were then intersected to identify liver and lung tumor-specific signatures by filtering out differentially expressed genes in the normal organs. Pathway enrichment analysis was performed on the resulting organ/tumor-specific gene sets using enrichR^73^, and the top enriched TFs were plotted using combined scores in bar plot format.

#### scRNA-seq analysis

Raw scRNA-seq data for ref. 47 were downloaded from the Gene Expression Omnibus (GSE155698). We kept cells using filters “min.cells > 100, nFeature_RNA > 500, nCount_RNA > 2500, percent.mt < 25”. SCTransform was used to regress out percent mitochondrial RNA, and nearest neighbors were found using FindNeighbors with dims = 1:30. Clusters were identified using resolution = 0.8, and cell types were annotated using R packages celldex, SingleR, Azimuth, and custom gene sets^85,86^. Only Ductal and acinar cells were retained for calculating gene signature scores from our mouse multiomic data using AddModuleScore from Seurat^86^. FeaturePlot was used to visualize the differential peak openings from our mouse LiverOPEN and LungOPEN ATAC signatures.

### Statistics and reproducibility

Statistical analyses were performed with GraphPad Prism (v.9), R (v4.3.1) and Python (v.3.6.4). Pooled data are presented as mean ± s.e.m. Sample size, error bars and statistical methods are reported in the figure legends. Exact p-values are shown in figures or associated legends. RNA- and ATAC-seq data were analyzed as described in the respective sections above. No statistical methods were used to predetermine sample size in the mouse studies. Mice were randomized into different treatment groups (for example, +Dox vs. –Dox). The investigators were not blinded to allocation during experiments and outcome assessment.

### Reporting summary

Further information on research design is available in the Nature Research Reporting Summary linked to this paper.

## DATA AVAILABILITY

All sequencing data will be deposited to GEO prior to publication. All other data supporting the findings of this study will be made available upon reasonable request.

## CODE AVAILABILITY

No unique code was developed for this study.

## ACKNOWLEDGEMENTS

We thank M. Chalarca, K. Rybczyk, and the MSKCC Research Animal Resource Center for technical assistance with mouse work; Z. Zhao, A. Kahn, Y. Zhao, S. Yang, S. Young, Y. Furuta, and the MSKCC Mouse Genetics Core for assistance with the generation of ESC-derived mouse models; R. Gardner and the MSKCC Flow Cytometry Core facility staff for assistance with cell sorting; O. Chaudhary and the MSKCC Single-cell Analytics Innovation Lab (SAIL) for technical assistance with scMultiomics; C. Burdziak and R. Sharma for advice on scMultiome analysis; F.J. Sánchez-Rivera, R. Mezzadra, J. P. Morris IV, J. Reyes and the rest of the Lowe laboratory for advice and helpful discussions; and J. Novak for manuscript editing. Mouse and human illustrations were created with BioRender (https://biorender.com/). We acknowledge Core funding by an NCI Cancer Center Support Grant (P30 CA08748), Cycle for Survival, and the Marie-Josée and Henry R. Kravis Center for Molecular Oncology. K.M.T. was supported by the Jane Coffin Childs Memorial Fund for Medical Research, a Shulamit Katzman Endowed Postdoctoral Research Fellowship, and an NIH K99/R00 Transition to Independence Award (5K99CA266939). F.M.B. was supported by a GMTEC Postdoctoral Fellowship, an MSKCC Translational Research Oncology Training Fellowship (5T32CA160001) and a Young Investigator Award by the Edward P. Evans Foundation. D.A.C. was supported by the Spanish Fundación Ramón Areces Postdoctoral Fellowship and is recipient of the La Caixa Postdoctoral Junior Leader Fellowship (LCF/BQ/PI20/11760006). G.L. was supported by an NIH F32 grant (1F32CA177072) and an American Cancer Society Fellowship (ACS PF-13-037-01-DMC). T.B. received support from the William C. and Joyce C. O’Neil Charitable Trust and the MSK Single Cell Sequencing Initiative. This work was supported by MSKCC’s David Rubenstein Center for Pancreatic Research Pilot Project (S.W.L.); the Alan and Sandra Gerry Metastasis and Tumor Ecosystems Center (J.M. and S.W.L.); the Lustgarten Foundation Research Investigator Award (S.W.L.); the Agilent Thought Leader Program (S.W.L.); and NIH grants P01CA13106 (S.W.L.) and R35CA252978 (J.M.). S.W.L. is the Geoffrey Beene Chair of Cancer Biology, D.P. is the Alan and Sandra Gerry Endowed Chair of Computational Biology, and J.M. is the Marie-Josée and Henry R. Kravis Chair. S.W.L. and D.P. are Investigators of the Howard Hughes Medical Institute.

## AUTHOR CONTRIBUTIONS

K.M.T. conceived the study, designed and performed experiments, analyzed data, and wrote the manuscript. F.M.B., D.A.C. and G.L. designed and performed experiments, and analyzed data. Y.-J.H. analyzed all bulk RNA-seq and scRNA/ATAC-seq data, including bulk RNA-ATAC-seq integration and analysis of public datasets. R.K. analyzed bulk ATAC-seq data. T.B. analyzed sparse WGS data. J.S., S.T., A.N.W. and W.L. assisted in experiments. J.E.W. helped interpret histopathology data. I.M. performed scRNA/ATAC-seq, with supervision and critical input from D.P. and R.C. N.D. assisted with data analysis. J.M., D.P. and C.A.I.-D. provided critical intellectual input and guidance in experimental design and data interpretation. S.W.L. conceived and supervised the study and wrote the manuscript. All authors read the manuscript.

## COMPETING INTERESTS

S.W.L. is a consultant for Fate Therapeutics, and is a consultant and holds equity in Blueprint Medicines, ORIC Pharmaceuticals, Mirimus, PMV Pharmaceuticals, Faeth Therapeutics, and Senescea Therapeutics. R.P.K. is a co-founder of and consultant for Econic Biosciences. D.P. is on the scientific advisory board of Insitro. J.M. holds equity in Scholar Rock. All other authors have no competing interests. None of these affiliations represent a conflict of interest with respect to the design or execution of this study or interpretation of data presented in this report.

## FIGURE LEGENDS

**Extended Data Figure 1.**
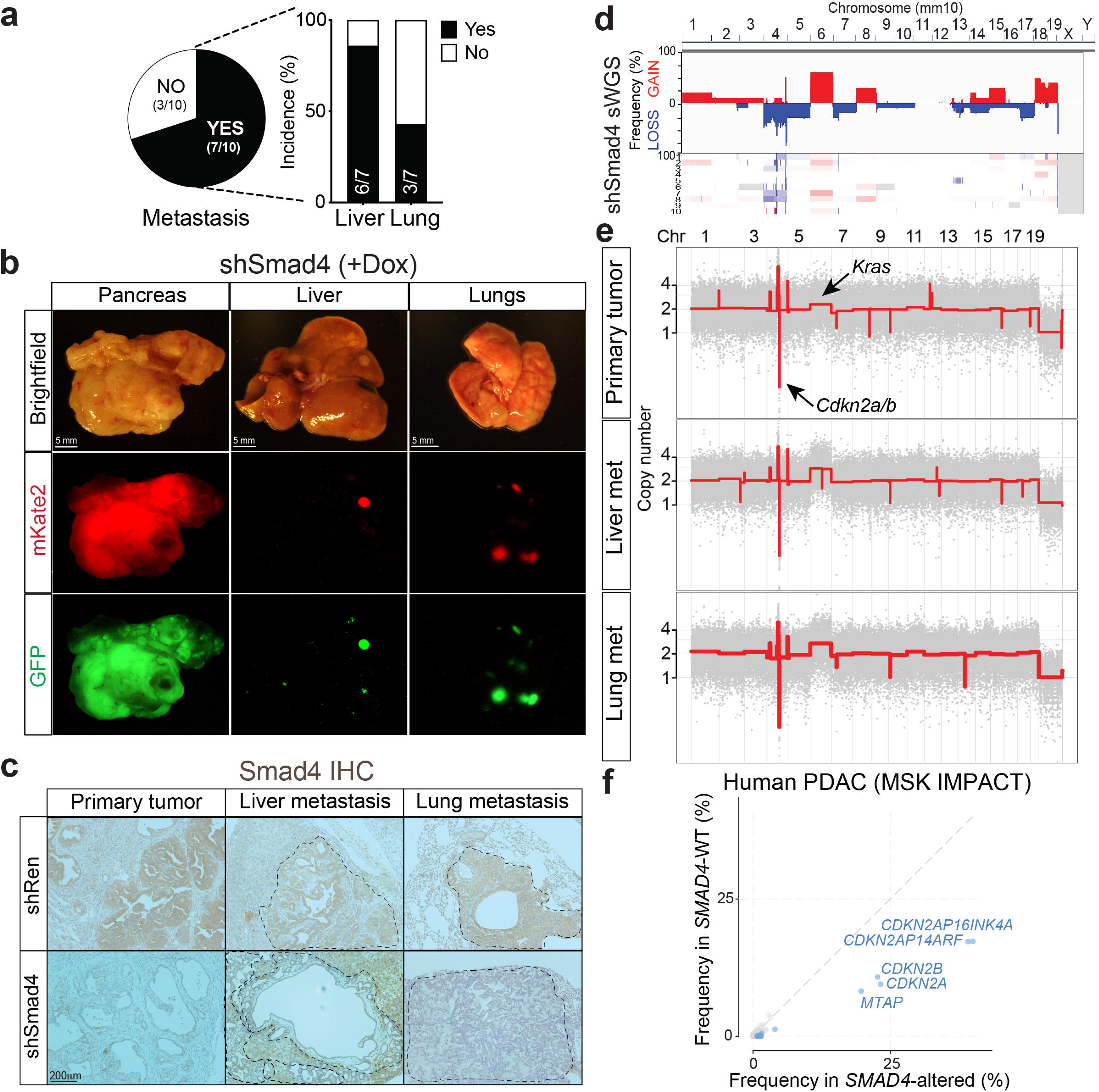
**(a)** Overall (pie chart) and organ-specific (stacked bar graph) frequency of metastasis in KC-shSmad4 mice. **(b)** Representative macroscopic images of tumor-bearing pancreas, liver, and lungs from KC-shSmad4 mice (+Dox) at endpoint. Fluorescent images match the corresponding brightfield images. Data are representative of 10 (pancreas), 6 (liver), and 3 (lungs) independent mice. **(c)** Representative immunohistochemistry (IHC) staining for SMAD4 in primary and metastatic KC-shSmad4 tumors (+Dox). Dashed lines demarcate metastases. Data are representative of 3 independent mice. **(d)** sWGS analysis of genome-wide copy number alterations in KC-shSmad4 primary tumor-derived cell lines (n=10 independent mice). Frequency plot is shown on the top and individual sample tracks are provided on the bottom. **(e)** Representative sWGS analysis of genome-wide copy number alterations in KC-shSmad4 tumor-derived cell lines from matched primary tumors, liver and lung metastases. The *Kras* and *Cdkn2a/b* loci are highlighted (arrows). Data are representative of 3 independent mice. **(f)** Frequency of the indicated homozygous deletions in *SMAD4*-altered (mutated or homozygously deleted, n=959) or wild-type (WT, n=3188) PDAC tumors in the MSK-IMPACT cohort (*CDKN2A/B*, p<10^-10^; q<10^-10^). *CDKN2A/B* and their adjacent gene *MTAP* are highlighted.

**Extended Data Figure 2.**
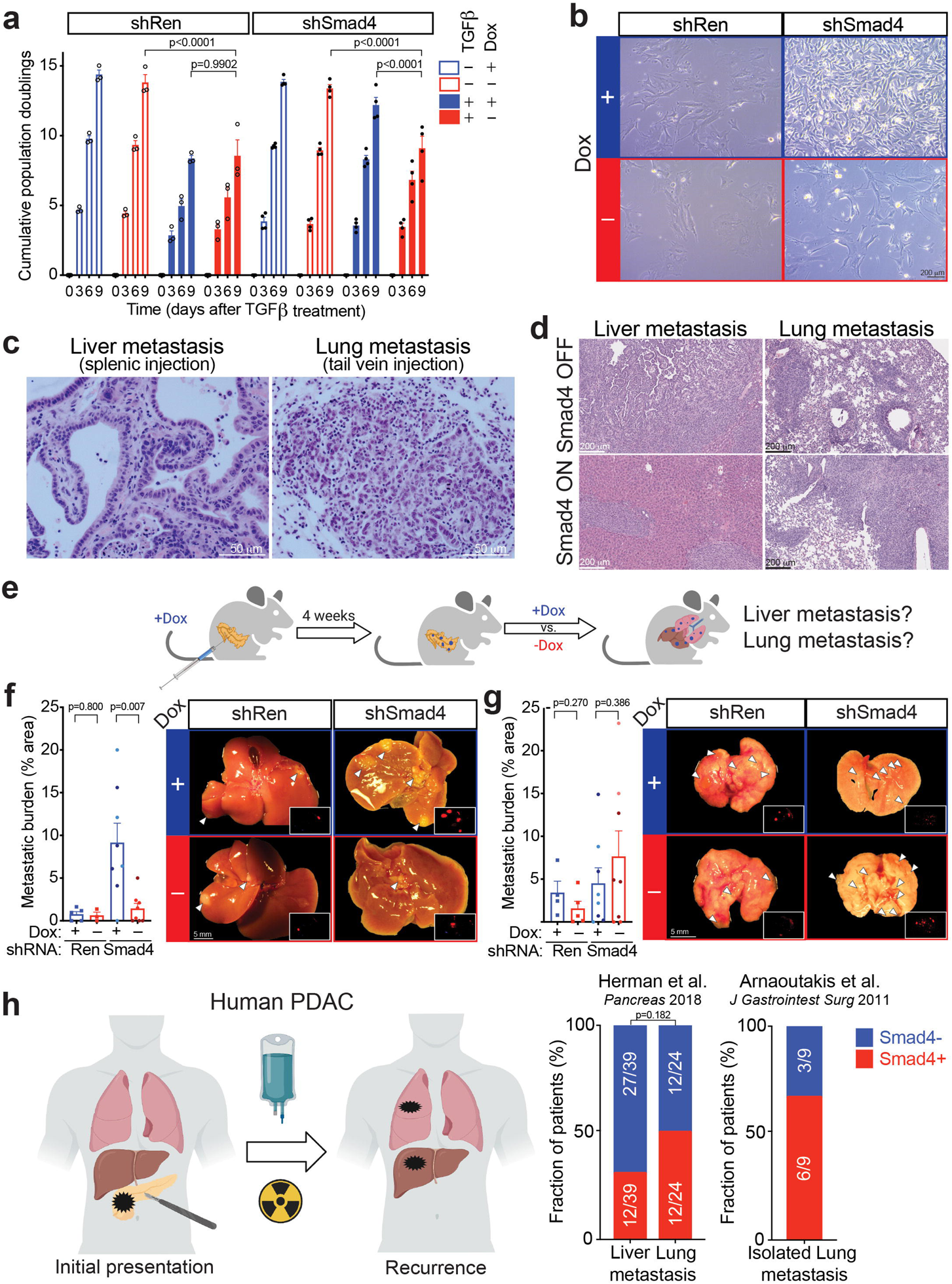
**(a)** Proliferation assays of KC-shRen and KC-shSmad4 tumor-derived cell lines +/-Dox +/-TGFβ *in vitro* (n=3 and 4 independently derived cell lines for KC-shRen and KC-shSmad4, respectively). **(b)** Representative brightfield images of KC-shRen and KC-shSmad4 cells after TGFβ treatment +/-Dox for 6 days. Data are representative of 3 independent cell lines per genotype. **(c)** Representative H&E staining of liver and lung metastases generated by intrasplenic or tail vein injections of KC-shSmad4 cell lines. Data are representative of 20 metastases across 4 independent mice per organ site. **(d)** Representative H&E staining of liver and lung metastases from KC-shSmad4 cells without (Smad4 OFF) or with *Smad4* restoration (Smad4 ON). Data are representative of 6 independent mice per organ site. **(e)** Schematic of orthotopic experiments with KC-shRen and KC-shSmad4 cells for analysis of metastasis burden. **(f)** Analysis of liver metastasis burden after orthotopic injections of KC-shRen or KC-shSmad4 cells with or without subsequent Dox withdrawal. (Left) Percent-area quantifications at endpoint (n=5, 3, 8, and 8 independent mice for groups shown, left to right). Different color shading indicates independent cell lines. (Right) Representative macroscopic images of tumor-bearing livers. Insets show tumor-constitutive mKate2 reporter. **(g)** Analysis of lung metastasis burden after orthotopic injections of KC-shRen or KC-shSmad4 cells with or without subsequent Dox withdrawal. (Left) Percent-area quantifications at endpoint (n=4, 5, 8, and 8 independent mice for groups shown, left to right). Different color shading indicates independent cell lines. (Right) Representative macroscopic images of tumor-bearing lungs. Insets show tumor-constitutive mKate2 reporter. **(h)** (Left) Schematic of metastasis recurrence studies in human PDAC^29,30^. (Right) Fraction of patients whose tumors stained positive or negative for SMAD4 by IHC, according to site of recurrence. Exact numbers are indicated on each bar Statistical analysis: (a) Two-way ANOVA; (f, g) Unpaired two-tailed t-test; (h) Fisher’s exact test.

**Extended Data Figure 3.**
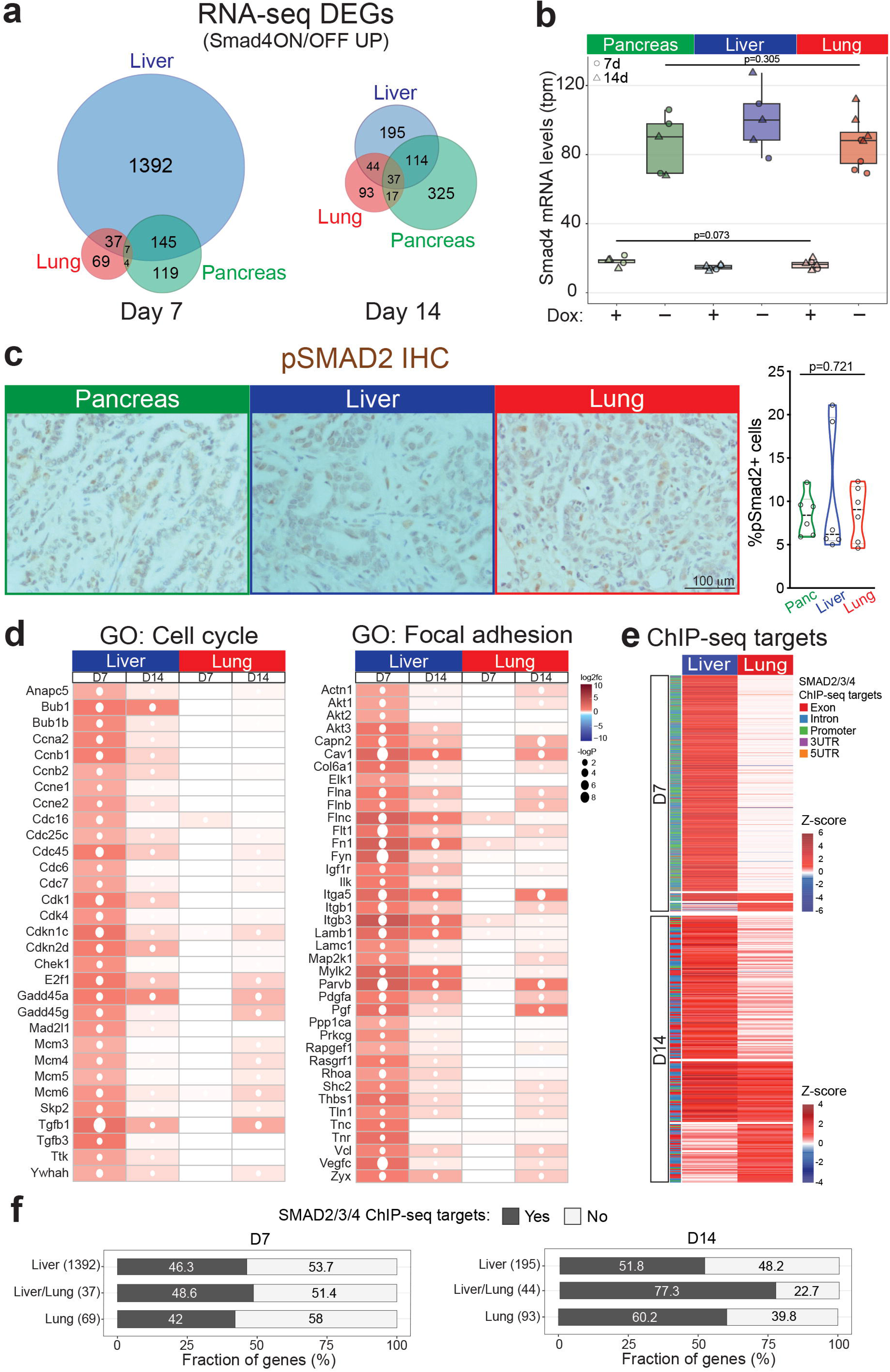
**(a)** Overlap between upregulated genes in tumor cells from the pancreas, liver, or lungs at 7 and 14 days after Dox withdrawal. Numbers reflect genes in each category. Complete gene lists are provided in ED Table 2. DEGs = <u>d</u>ifferentially <u>e</u>xpressed <u>g</u>ene<u>s</u>. **(b)** Normalized *Smad4* mRNA levels (RNA-seq data; tpm = transcripts per million) in KC-shSmad4 tumor cells from the pancreas, liver, or lungs +/-Dox for 7 or 14 days. **(c)** Representative IHC staining for pSMAD2 in KC-shSmad4 tumors (+Dox) from the pancreas, liver, or lungs. Quantifications (%pSMAD2 cells) are shown on the right (n=6 independent metastases from two mice). Statistical analysis by one-way ANOVA. **(d)** Heatmap of differentially expressed genes from the indicated KEGG pathways. Average log2 fold-change and p-values are shown for each organ and timepoint. **(e)** Heatmap of SMAD4-dependent SMAD2/3 ChIP target genes that are differentially upregulated in liver vs. lung metastases at 7 or 14 days after Dox withdrawal. Average RNA-seq log_2_ fold-change values are shown for Smad4 ON vs. OFF in each organ and timepoint. The type of genomic binding region is color-annotated on the left. Complete gene lists are provided in ED Table 2. **(f)** Fraction of upregulated genes in liver and/or lung metastases at 7 or 14 days of Dox withdrawal that are SMAD4-dependent SMAD2/3 ChIP-seq targets. Absolute number of genes per group is shown in parentheses.

**Extended Data Figure 4.**
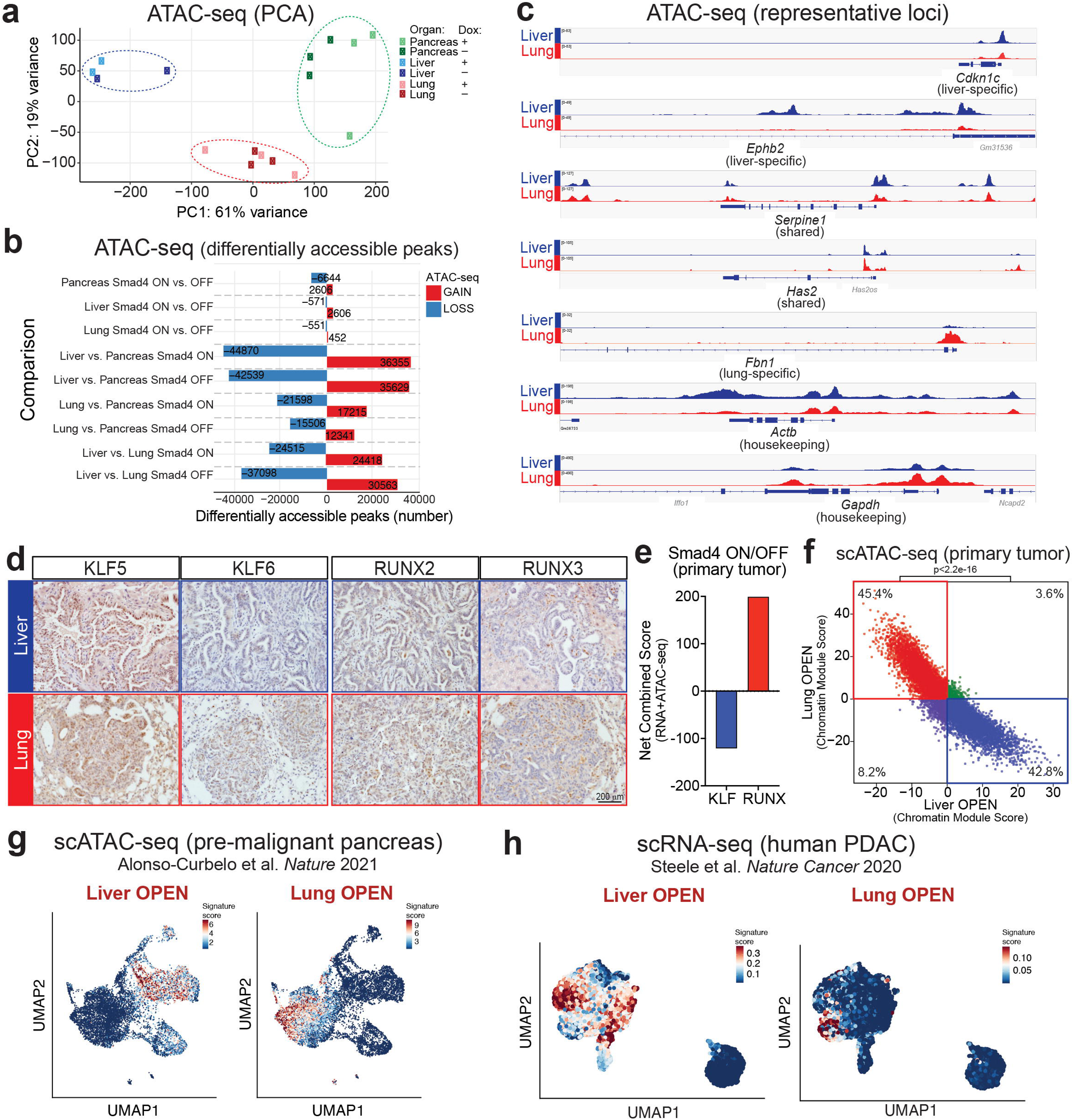
**(a)** Principal component analysis (PCA) of ATAC-seq data from tumor (mKate2^+^) cells isolated from the pancreas, liver, or lungs. Analysis is based on peak normalization. Each sample corresponds to an independent mouse. Circled areas highlight separation based on organ site. **(b)** Number of statistically significant differentially accessible ATAC-seq peaks. Exact number of gained/lost peaks are shown for each comparison (absolute fold change ≥ 1.5; FDR ≤0.1). **(c)** Representative ATAC-seq tracks at loci with liver-specific, lung-specific, or liver/lung-shared chromatin opening in KC-shSmad4 tumor cells (mKate2^+^GFP^+^). Housekeeping genes are shown as a reference. Data are representative of 3 independent tumors. y-axis scale range is indicated per lane as normalized read counts. **(d)** Representative IHC staining for the indicated TFs in KC-shSmad4 (+Dox) liver or lung metastases. Data are representative of 20 metastases across 4 independent mice. **(e)** Net change in ATAC-RNA combined scores for the KLF and RUNX TF families in Smad4 ON vs. OFF primary tumors at day 14. This metric infers the probability that a given TF family is impacted by Smad4 restoration, based on a combined change in ATAC-seq motif accessibility and RNA-seq levels of its predicted target genes (see Methods for details). **(f)** Mutual exclusivity of liver- and lung-specific open chromatin signatures in primary tumors. The plot shows the distribution of cells from KC-shSmad4 (+Dox) primary tumors according to their enrichment of liver-(LiverOPEN) and lung-specific (LungOPEN) open chromatin signatures derived from bulk ATAC-seq (see Methods for details). Each dot corresponds to an independent cell. The proportions of cells with liver-only (blue) and lung-only (red) signatures were compared to those with both liver and lung signatures (green) and those with neither signature (purple) using Chi-squared test. **(g)** UMAP visualization of scATAC-seq profiles of *Kras*-mutant pancreatic epithelial cells (mKate2^+^) *+/-* cerulein injury (n=1 mouse for each) as described in ref. 45 (see Methods for details). Signature scores based on liver- or lung-specific ATAC-open peaks from bulk ATAC-seq data are displayed in color per individual cell. **(h)** UMAP visualization of scRNA-seq profiles of human primary PDAC tumor cells from ref. 47 (see Methods for details). Signature scores based on scRNA-seq profiles of mouse liver- or lung-specific ATAC-open cell populations (from the matching scATAC-seq multiomics data) are displayed in color per individual cell.

**Extended Data Figure 5.**
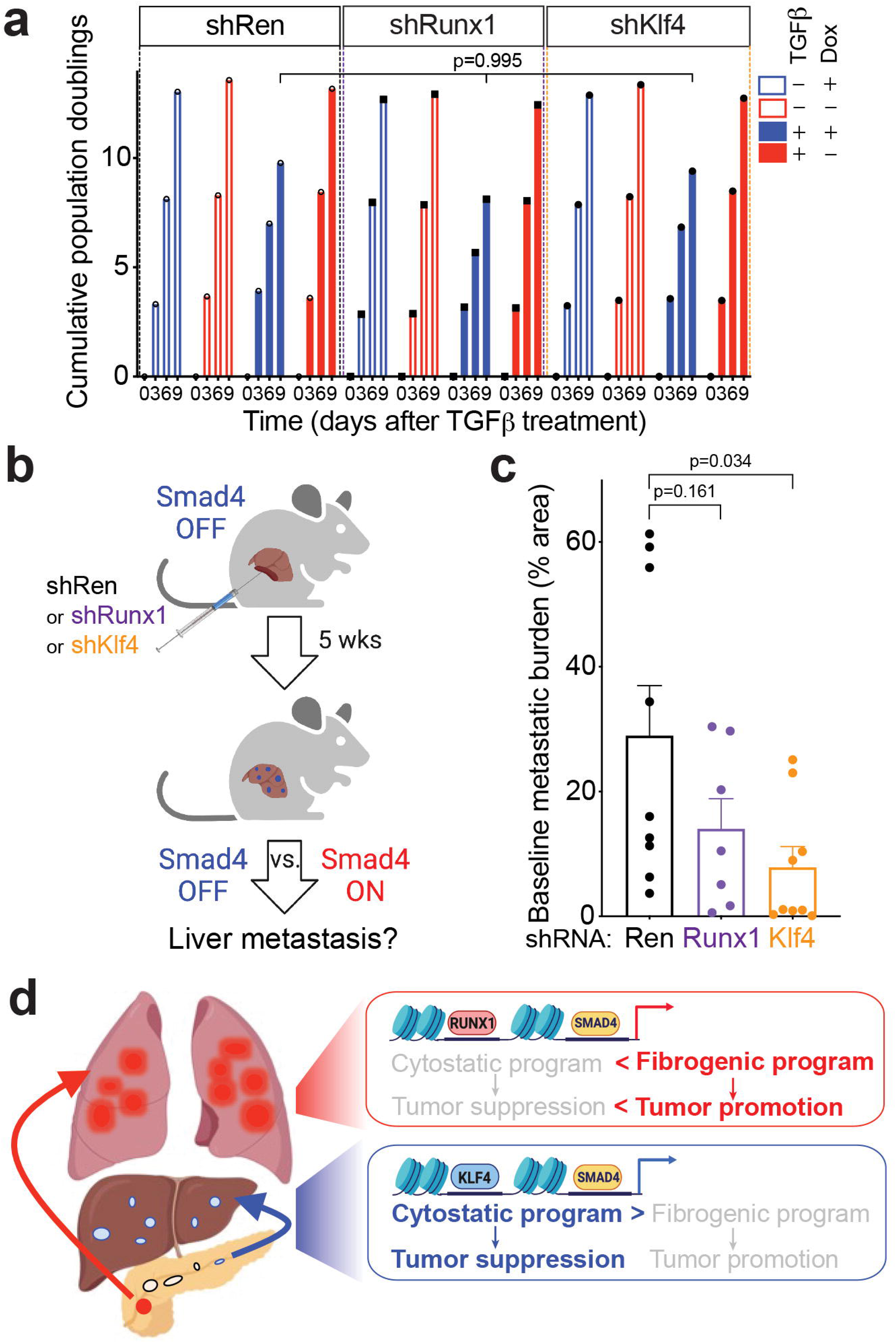
**(a)** Proliferation of KC-shSmad4 cell lines harboring the indicated stable shRNAs +/-Dox and +/- TGFβ *in vitro* (average of three technical replicates). Statistical analysis by one-way ANOVA. **(b)** Schematic of intrasplenic experiments with KC-shSmad4 cells harboring TF knockdowns. **(c)** Quantifications of baseline metastasis burden (% tumor area) after intrasplenic injections of KC-shSmad4 cells under continuous Dox administration (n=8, 7, and 9 independent mice per group, respectively). Livers were harvested 64-66 days after injection. Statistical analysis by unpaired two-tailed t-test. **(d)** Model of the organ-specific interplay between *SMAD4* and KLF4/RUNX1-associated chromatin states. In the liver, active KLF4 allows SMAD4 to act at genes that promote cytostasis, thus necessitating the inactivation of *Smad4*. In the lungs, more speculatively, active RUNX1 allows SMAD4 to act at pro-fibrotic genes unopposed by its cytostatic program.

